# Interference hypothesis for recombination suppression in chromosomal inversion heterozygotes: A formal genetics analysis in *Drosophila melanogaster*

**DOI:** 10.1101/305490

**Authors:** Spencer A. Koury

**Affiliations:** Department of Ecology and Evolution Stony Brook University Stony Brook, NY 11794

**Keywords:** Chromosomal Inversion, Recombination Suppression, Crossover Interference

## Abstract

Suppressed recombination in chromosomal inversion heterozygotes is a well-known but poorly understood phenomenon. Recombination suppression is the result of at least four different regional effects of inversion heterozygosity; surprisingly, this includes areas outside inversions where there are no barriers to normal pairing, synapsis, double strand break formation, or recovery of crossover products. The interference hypothesis of recombination suppression proposes heterozygous inversion breakpoints possess chiasma-like properties such that recombination suppression extends from breakpoints in a process analogous to crossover interference. This hypothesis is qualitatively consistent with chromosome-wide patterns of recombination suppression, unifying the various regional effects under a single mechanism. The present study generates quantitative predictions for this hypothesis using a probabilistic model of crossover interference as a stationary renewal point process with gamma-distributed interarrival distances. These predictions were tested with formal genetic data (>40,000 meioses) on crossing-over in intervals external and adjacent to four cosmopolitan inversions of *Drosophila melanogaster*. The decay of recombination suppression differs for each of the four cosmopolitan inversions, with a strong dependence on proximity to the centromere. Counterintuitively, greater than expected recombination was observed in centromeric regions for the most proximally placed inversions. Structural features of chromosomes are discussed as potentially confounding variables in modelling recombination suppression. The true form of the decay function is not currently known and will require high-resolution mapping of rare crossover events in regions immediately adjacent to inversion breakpoints. A simple extension of the present experimental system can be conducted as a genetic screen for those rare crossover events.

## INTRODUCTION

Based solely on formal genetic mapping data Alfred Sturtevant predicted the existence of chromosomal inversions. This hypothesis was based on four discoveries also made by Sturtevant; first, crossover data could be used to create genetic maps of linear chromosomes (Sturtevant 1913), second, chromosomes segregating in *Drosophila melanogaster* populations carried crossover modifiers (Sturtevant 1917), third, complementation tests show *D. simulans* and *D. melanogaster* share genes (Sturtevant 1920), and fourth, those shared genes were not necessarily in the same order (Sturtevant 1921). These experiments suggested the elegant hypothesis that the polymorphic crossover modifiers in *D. melanogaster* were in fact inverted gene orders. Cytological confirmation of this farsighted hypothesis came over a decade later (Painter 1933), and the evolutionary analysis of inversion polymorphism quickly followed, again pioneered by Sturtevant (Sturtevant and Dobzhansky 1936; Sturtevant and Mather 1938; Sturtevant and Novitski 1941). Although scientific consensus is not complete, it is likely that the evolutionary persistence of inversions is either a direct or indirect effect of their ability to suppress recombination in the heterozygous state (Kirkpatrick and Barton 2006; Hoffmann and Rieseberg 2008). Recombination suppression was Sturtevant’s first clue to the existence of inversions, but more than a century later the dimensions of this complex meiotic phenotype are still poorly understood. In the present paper, the reasons for an incomplete understanding of the fundamental property of inversions are briefly summarized and then an interference model of recombination suppression is introduced as a potentially unifying hypothesis.

In *D. melanogaster* recombination suppression is the combined effect of at least four phenomena which occur in several distinct regions relative to an inversion breakpoint (figure 1A). Suppression effects have been extensively documented, but the use of heterogeneous live material (in terms of genetic backgrounds, inversions, markers, and species) to study reduction in recombination rates complicates integration and interpretation of a sizeable literature (Krimbas and Powell 1992). Classical work focused on structural features like size and position of inverted regions as the major determinants of recombination suppression effects, but recent experimental population genetic research has demonstrated there is substantial genic variation for the genomic rate and chromosomal distribution of recombination events segregating in many species (Bauer *et al*. 2013; Kong *et al*. 2014; Johnston *et al*. 2016), including *D. melanogaster* (Comeron *et al*. 2012; Hunter *et al*. 2016). It is currently unknown to what degree an inversion’s recombination profile is a function of its structural features (size and position of inverted regions), as opposed to the genetic elements that are in strong linkage disequilibrium with inverted regions. However, several generalizations do apply to this diverse set of experiments; first, the strongest effects of recombination suppression are experienced in the local vicinity of inversions and second, diffuse recombination effects are experienced throughout the genome, even in physically unlinked regions (Krimbas and Powell 1992).

**Figure 1.**
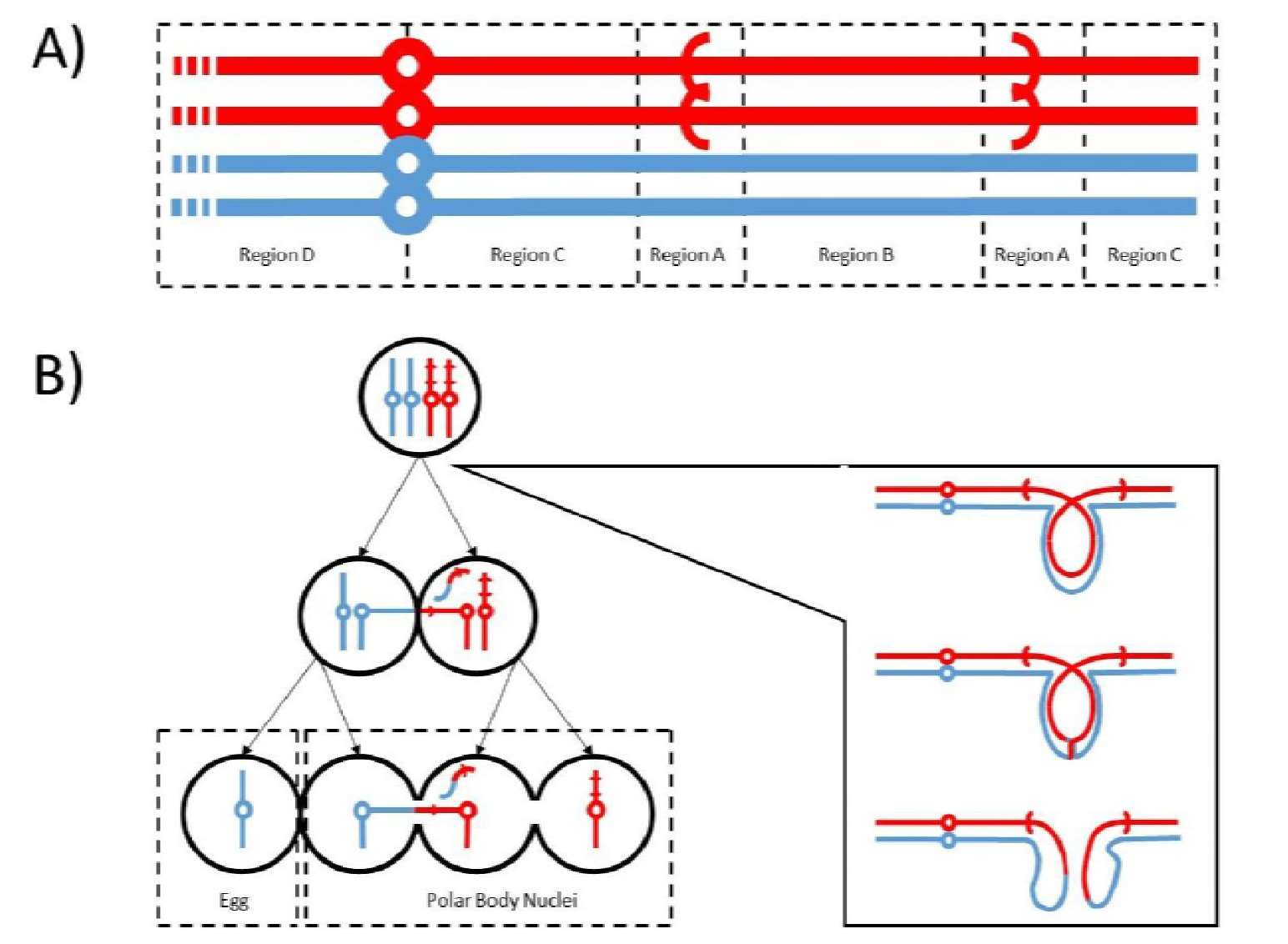
A) Chromosomes of an inversion heterozygote as a four strand bundle highlighting the regions of different recombination suppression effects relative to inversion breakpoints (indicated by parentheses). B) The formation of acentric and dicentric chromosomes as a consequence of crossing-over in the inversion loop, illustrated to the right, and selective elimination of dicentric chromosomes by confinement to internal positions and therefore polar bodies in the linear array of meiotic products.

In the region immediately adjacent to inversion breakpoints (figure 1A region A, at least 2 Mb in *D. pseudoobscura* (Stevison *et al*. 2011)), crossing-over has not been observed and thus recombination suppression is assumed to be complete. Historically researchers thought failure of homologous chromosomes to synapse in region A caused this complete suppression (Dobzhansky 1931; Sturtevant and Beadle 1936; Novitski and Braver 1954). More recently, cytological analysis of multiply inverted *D. melanogaster* X chromosomes revealed that pairing, synapsis, and the formation of double strand breaks are approximately normal (Gong *et al*. 2005). The weak effect of failed pairing or synapse immediately at inversion breakpoints is insufficient to explain complete suppression of recombination in region A. Gong *et al.’s* (2005) findings indicate crossovers are initiated at normal rates, although they are not resolved at normal rates. Thus, the mechanism by which complete suppression in region A occurs is unknown.

The most well-known effect of inversion heterozygosity is recombination suppression in the region internal to inversion breakpoints (figure 1A region B). This is because a single crossover event generates acentric and dicentric recombinant products that are never included in the functional egg *via* a meiotic drive mechanism unique to asymmetric meiosis in females and only known from studies in *Diptera* (figure 1B) (Sturtevant and Beadle 1936; McClintock 1941; Carson 1946; Madan *et al*. 1984). Recombination can and does occur in region B as demonstrated by the recovery of double crossover events under normal conditions and the ability to recover single crossover events under special experimental conditions (*e.g*., using compound X chromosomes) (Sturtevant and Beadle 1936). Using experimental constructs to avoid the mechanical difficulties associated with recombinants, or using viability reduction with pericentric inversions as an indirect measure, the recombination rate in region B is estimated to be about 25% of wildtype (Novitski and Braver 1954; Coyne *et al*. 1993; Navarro and Ruiz 1997). Despite the clear understanding of the selective elimination of aberrant products from single crossover events (figure 1B), the mechanism for the ~75% partial suppression of crossing-over in region B remains unknown.

In the regions external to inversion breakpoints (figure 1A regions C and D) recombination occurs, but generates a genetic map strongly influenced by proximity to inversion breakpoints. In region C there are no apparent barriers to normal pairing, crossing-over, and recovery of all meiotic products; nonetheless, recombination is still reduced in areas near the inversion (Sturtevant 1931; Sturtevant and Beadle 1936; Grell 1962). In regions very distant from inversion breakpoints (figure 1A region D) recombination has sometimes been observed to increase and this is termed the intrachromosomal effect (Dobzhansky and Sturtevant 1931; Dobzhansky 1933). A similar observation of increased recombination on non-homologous chromosomes is termed the interchromosomal effect (Schultz and Redfield 1951). The mechanisms for the partial suppression in region C, the intrachromosomal effect in region D, and interchromosomal effect on other chromosomes is unknown (Lucchesi and Suzuki 1968). Further complicating matters, the relationship amongst, and the transition between the mechanisms responsible for the complete, partial, and negative suppression (regions A, C, and D respectively) remains obscure.

Clearly, recombination suppression *sensu lato* is a complex phenotype consisting of several types of crossover modification at different distances from inversion breakpoints which vary in both magnitude and extent. Most of these effects were described in the pre-molecular era of biology. The application of molecular genetics methods in the multiply inverted chromosome of *D. melanogaster* has shown older assumptions about pairing, synapsis, and formation of double strand breaks near inversion breakpoints to be incorrect (Gong *et al*. 2005; Miller *et al*. 2016a; Miller *et al*. 2016b). To explain the lack of recombination despite normal rates of crossover initiation Gong *et al*. (2005) suggest heterozygous inversion breakpoints possess the interference-like properties of chiasmata. The present paper provides a mathematical elaboration of the interference hypothesis for recombination suppression that has the potential to unify the disparate regional effects of inversion heterozygosity under a single mechanism.

Similar to recombination suppression, interference is a well-described phenomenon whose causal basis remains poorly understood (Speed 1996; Miller *et al*. 2016c). Interference is the non-independence of crossover events and is traditionally described by a coefficient of coincidence. Formally, interference is the conditional probability of crossing-over in a focal interval, given a crossover event elsewhere in the genome, scaled by the unconditional probability of crossing-over for the focal interval. Practically, quantification of interference requires multi-locus data and the coefficient of coincidence is expressed as the ratio of observed double crossovers to the expected number under crossover independence (*i.e.*, joint probability / product of marginal probabilities) (Muller 1916; Bailey 1961).

Interference was first observed in *D. melanogaster* (Sturtevant 1915; Muller 1916), and much of the mathematical development of interference theory has been based on single spore data generated from the *D. melanogaster* model system (Haldane 1919; Kosambi 1943; Bailey 1961). Modelling the underlying point process for the gold standard multi-locus datasets (Weinstein 1936; Morgan *et al*. 1938), multiple groups have all arrived at the same conclusion: interference in *D. melanogaster* is most accurately modeled as a stationary renewal process where interarrival times follow a gamma distribution with shape parameter (*γ*) of approximately 5 and a rate parameter (*μ*) scaled to cumulative interval distances (Foss *et al*. 1993; McPeek and Speed 1995; Zhao *et al*. 1995). When the shape parameter is an integer, the counting model of Foss *et al*. (1993), the chi-square model of Zhao *et al*. (1995), and gamma model of McPeek and Speed (1995) are equivalent. Following McPeek and Speed (1995), the coefficient of coincidence (*C*) as a function of genetic distance (*z*) from the previous crossover event is

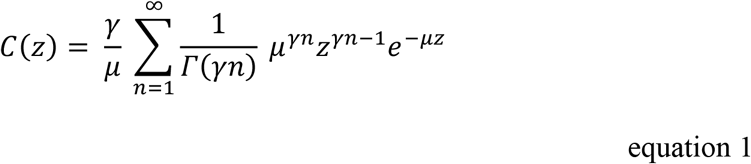

where *γ* is the shape parameter, *μ* is the rate parameter, and *n* is an auxiliary variable. The *D. melanogaster* specific parameterization of the coincidence function above suggests negligible recombination (<0.0001) will occur within 5 map units of inversion breakpoints, recombination suppression will decay over approximately 35 map units, and a weak elevation of recombination rates above wildtype will be observed in regions 40-60 map units distant from the inversion breakpoints (where map units refers to cM for the standard arrangement genetic map of *D. melanogaster*). These predictions are qualitatively consistent with the effects described for the regions A, C, and D in *D. melanogaster* (Sturtevant 1931; Sturtevant and Beadle 1936; Grell 1962; Miller *et al*. 2016b)

The interference model also allows quantitative prediction of the magnitude and spatial extent of recombination suppression for any inversion, and is graphically illustrated for a hypothetical case in figure 2. The predictions are generated by using the McPeek and Speed (1995) stationary renewal model of interference, where probability density of interarrival distances are gamma distributed with shape parameter *γ* = *5* and intensity parameter *μ* = *2γd* = *10*, and where *d* is the cumulative interval distances. This intensity parameter reflects the expected number of crossovers for the whole chromosome, *d* = *1* for a chromosome approximately 1 Morgan in length, such that it encompasses the absolute suppression of the inverted region and any intrachromosomal effects. To adapt this model to the present study of recombination suppression, heterozygous inversion breakpoints were treated as chiasmata and suppression effects were allowed to extend across the centromere. With these assumptions the predicted recombination fraction in inversion heterozygotes is equal to the area under the curve for the coincidence function (equation 1, figure 2).

**Figure 2.**
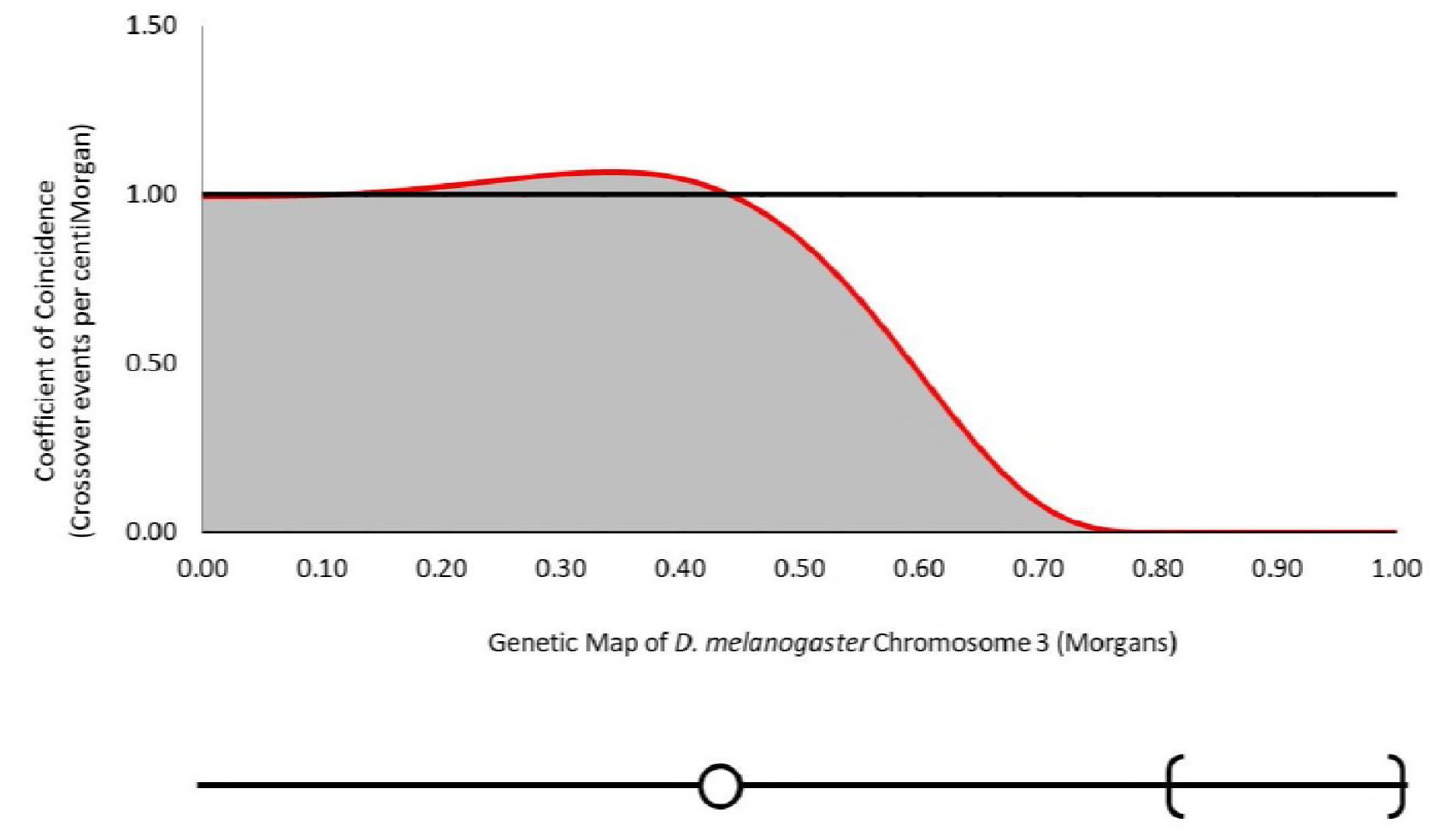
The coincidence function for a hypothetical terminal inversion based on the gamma model of (McPeek and Speed 1995). The black line represents unconditional probability of crossing-over for homokaryotypes and the red line is the conditional probability under the interference hypothesis. The expected recombination fraction for a heterokaryotype is equal to the gray shaded area under the curve.

The present study characterizes the recombination suppression effects of heterokaryotypes for four different cosmopolitan paracentric inversions of Muller element E in *D. melanogaster*. Formal genetic data is presented on recombination fractions for two adjacent chromosomal intervals flanked by dominant phenotypic markers. The balanced design of experiments incorporating both markers and inversions in all possible *cis* and *trans* arrangements allows the simultaneous estimation of viability effects and the strength of recombination suppression (Bailey 1961). This is a critical aspect of experimental design as both viability and suppression effects in each of the five gene arrangements differed in statistical significance. A *χ^2^* goodness-of-fit test was applied to the viability-corrected recombination fractions to evaluate the interference hypothesis. Viability-corrected counts of crossover events did not fit the expectations from the interference model, with the counterintuitive result that the decay of recombination suppression is faster for inversions closer to the centromere (*i.e*. recombination rate are *higher*, not lower, than expected near the centromere). The description of position dependence of interference effects identifies the necessary theoretical developments and experimental controls for future tests of the interference hypothesis of recombination suppression. Finally, a simple extension of the current experimental system to conduct genetic screens for those rare crossover events necessary for fitting the suppression decay function is described.

## MATERIALS AND METHODS

### Stock Construction

Inbred lines carrying the standard arrangement (*St*) and cosmopolitan inversions *In(3R)C, In(3R)K, In(3R)Mo*, and *In(3R)P* were drawn from the *Drosophila melanogaster* Genetic Reference Panel (DGRP) (table 1) (figure 3). Inversions were identified by polytene chromosome squashes of third instar larva salivary glands and confirmed with PCR amplification of inversion breakpoints where possible. The inversion *In(3R)C* was found in a single DGRP stock only (*DGRP 907*). In this stock *In(3R)C* was maintained in the heterozygous state (despite 20 generation of full sib mating) due to recessive viability and fertility effects segregating in the line (Koury Unpublished). All other inversions were selected from lines free of heterogenic regions, residual heterozygosity, or major fitness defects (Langley *et al*. 2012; Mackay *et al*. 2012; Koury Unpublished).

**Table 1.**
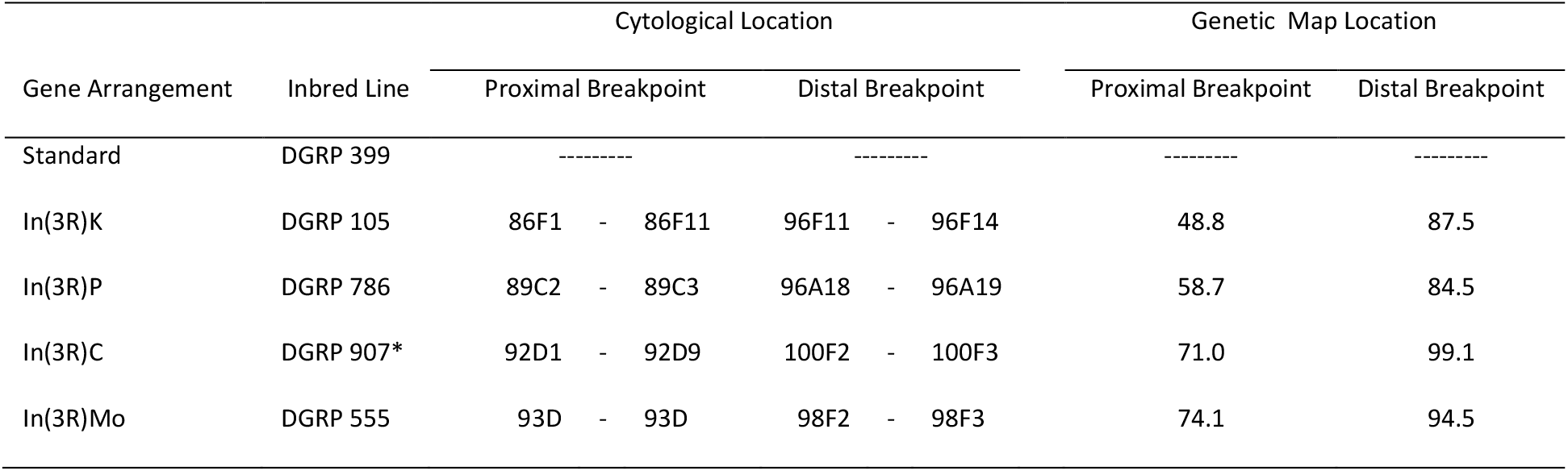
Cytological and genetic map location of gene arrangements used in this study.

**Figure 3.**
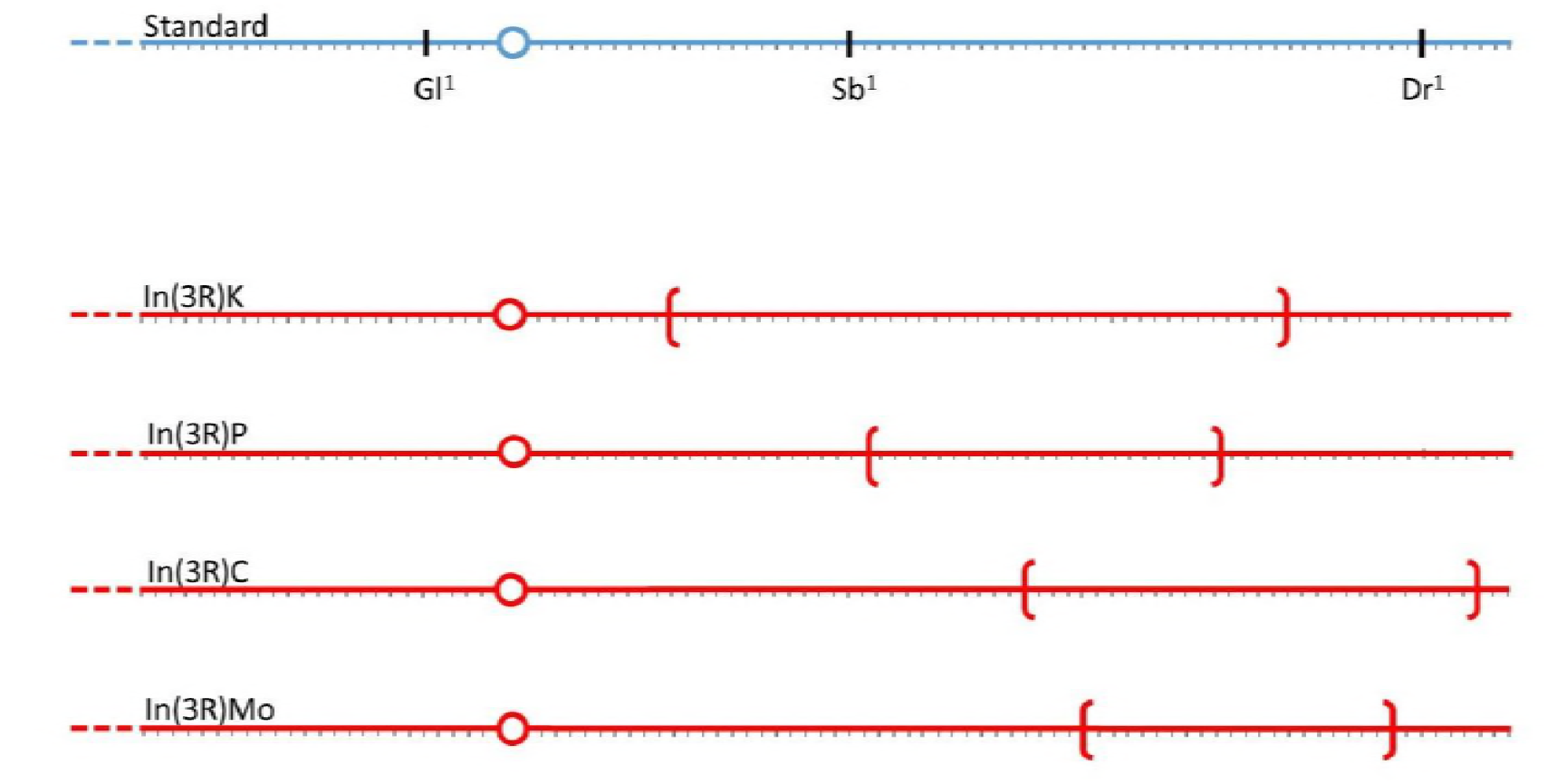
The relative position of phenotypic markers and the four cosmopolitan inversions of *D. melanogaster* Muller element E. The centromere is drawn as a circle and inversion breakpoints are denoted by parentheses. Diagram is to scaled based on the standard genetic map. For cytological and genetic map locations see table 1.

Focal chromosomes were isolated by balancer chromosome assisted extraction and placed on a common, standard arrangement genetic background for the X, Y, mitochondrial, and second chromosome. The non-recombining heterochromatin-rich fourth chromosome underwent repeated backcrossing to the standard genetic background giving a >75% chance of full background replacement, a probability that increases with each generation subsequent to stock formation. The necessary crossing scheme required introduction of balancers and a marked translocation into the standard genetic background (*DGRP 399*) and is illustrated in supplemental figure 1. This procedure was repeated for all five third chromosome gene arrangements.

Three dominant phenotypic markers (*Gl^1^, Sb^1^*, and *Dr^1^*) were selected for this study because of their relative position to inversion breakpoints (figure 3) (Locus *Gl* has recently been renamed *DCTN1-p150* in Flybase). Notably, the three mutations causing dominant phenotypic effects are also recessive lethals. These mutations were recombined onto standard and inverted arrangements followed by repeated backcrossing for a minimum of ten generations after confirmation of inversion presence by chromosome squash and PCR. Two exceptions were made, *Dr^1^* on *In(3R)Mo* and *Sb^1^* on *In(3R)P*, because the marker loci were experimentally determined to exist in a region of near complete recombination suppression, as >10,000 meioses failed to produce desired recombinants (Koury unpublished).

All three dominant markers (*Gl^1^, Sb^1^*, and *Dr^1^*) were also introgressed into the common tester stock *Canton-S* (Bloomington Drosophila Stock Center #1) which has the standard arrangement on all chromosome arms. Sometime prior to 2011, *Canton-S* strain was contaminated at the Bloomington Drosophila Stock Center with a *y^-^w^-^* transgene that is also present in this study. *Canton-S* contributes equally to all experimental crosses and thus any defects, if present, cannot explain variation observed among these experiments. Finally, the isogenic stock *w*; 6326; 6326* was used for outcrossing the *F_1_* experimental females. A full list of stocks used in experiments is provided in supplemental table 1.

### Crossing Design

To generate *F_1_* experimental genotypes, three virgin females of genotype *Canton-S* were crossed to three males homozygous for a given gene arrangement *Standard, In(3R)C, In(3R)Mo, In(3R)P*, or *In(3R)K*, and hereafter collectively referred to as *In(3R)x*. Desired *F_1_* experimental genotypes were selected based on the presence of dominant phenotypic markers and outcrossed to male *w*; 6326; 6326*. The progeny of this cross (*F_2_*) were scored for recombination *via* dominant markers and non-disjunction *via* white eyed patroclinous exceptions (figure 4a).

**Figure 4.**
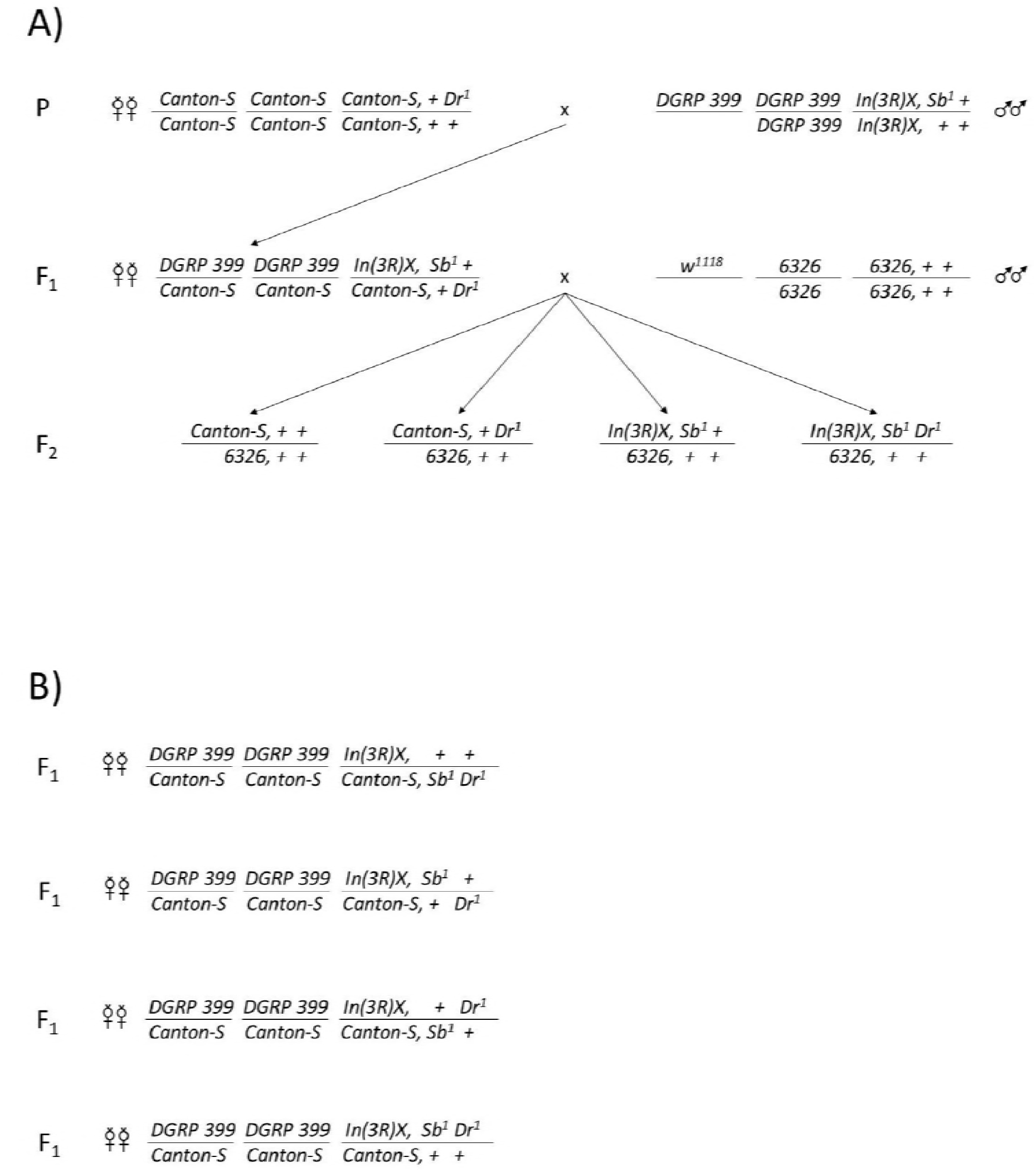
A) General crossing scheme of this experiment. B) The four alternative *F_1_* experimental genotypes used to perform both cis-trans and marker switching experiments.

A balanced design was employed to estimate genetic distances and simultaneously control for viability effects of dominant phenotypic markers, as well as direct fitness effects of the genetic backgrounds and inversions themselves. This balanced design was achieved by conducting both “marker switching” and “*cis-trans*” recombination experiments with all possible marker-inversion combinations on a common genetic background. Four different crosses were performed to generate *F_1_* females with all markers and inversions in a full factorial array. In the *P* generation, virgin *Canton-S* females with marker genotypes *Sb*^+^ *Dr*^+^, *Sb^1^ Dr*^+^, *Sb*^+^ *Dr^1^*, or *Sb^1^ Dr^1^* were mated with males homokaryotypic for one of the five gene arrangements carrying either *Sb^1^ Dr^1^, Sb*^+^ *Dr^1^, Sb^1^ Dr*^+^, or *Sb*^+^ *Dr*^+^, respectively. Thus, the females selected from the progeny as the *F_1_* experimental genotypes carried *Dr^1^, Sb^1^*, and *In(3R)x*, all in heterozygous state and in all possible linkage arrays on a common genetic background (figure 4B). For each gene arrangement a second experiment was conducted independently following the same methods, but using *Gl^1^, Sb^1^*, and *In(3R)x* in all possible combinations.

### Experimental Conditions

Experimental conditions follow the standard methods for mapping established by Bridges and Brehme (1944). To generate the *F_1_* experimental genotype three virgin females with the background *Canton-S* were crossed to three males for a given gene arrangement *In(3R)x*. Virgins of the desired *F_1_* genotype were collected over a three day period, aged an additional three days, then outcrossed to isogenic stock *6326*, which had the standard arrangement on all chromosome arms and the X-linked mutation *w^1118^*. Five *F_1_* experimental genotype virgin females were crossed with five *6326* males using light CO_2_ anesthesia, and after allowing 24 hours for recovery the mated group of ten individuals were tap transferred into half pint bottles with 30-40 ml of standard cornmeal-agar *Drosophila* food. Three replicate bottles were set for each cross. After five days of egg laying the *F_1_* adults were removed from bottles. A 2.5 inch × 2.5 inch blotting paper square was added to provide ample pupation sites with 0.05% v/v propionic acid added as needed to hydrate food. Emerging progeny (*F_2_*) were then scored daily for recombination (*via* dominant markers) and non-disjunction (*via* white-eyed patroclinous exceptions) for 15 days after the last eggs were laid (20 days after first eggs were laid). All vials and bottles were held at 25° C, greater than 50% relative humidity, under 24 hour light in a Percival incubator.

### Statistical Analysis

The crossing scheme outlined above yields a 2 × 2 × 2 full factorial design with respect to markers and inversions. With knowledge of the initial *P* genotypes, the frequency of each progeny phenotypic class can be further decomposed into the effect of recombination in addition to the viability effects of the dominant phenotypic markers and genetic background (*i.e*., 2 × 2 × 2 × 2 full factorial design). The angular transformation 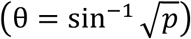 was applied to the observed frequency of each phenotypic class to fit ANOVA’s error term normality assumption (Sokal and Rohlf 1995). Data for each arrangement and was independently fit to the following model:

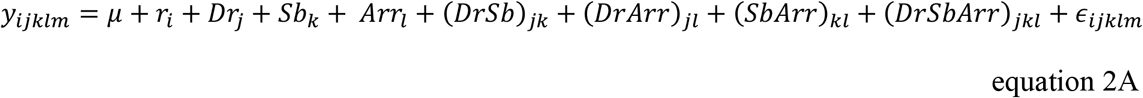

Where *y_ijklm_* is the angular transformed frequency of a phenotypic class, *r_i_* denotes whether the class is recombinant or nonrecombinant, *Dr_j_* + *Sb_k_* + *Arr_l_* are effects of markers *Dr^1^, Sb^1^* and *In(3R)x* for that phenotypic class. The analogous model for the *Gl* - *Sb* interval is:

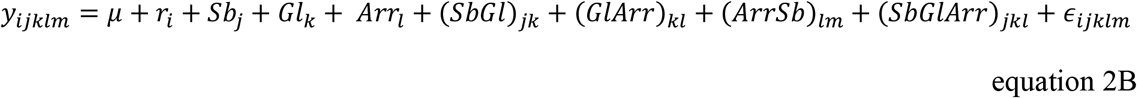

Thus for each of the 102 bottles in this study there were four data points each corresponding to a count for a unique phenotypic class. Each count was converted into a proportion and arcsine square root transformed for an analysis of variance corresponding to the linear model described above.

Effects of markers and gene arrangements were reported as a proportional viability of wildtype or the standard gene arrangement of *Canton-S*. Using the angular back-transformation (*p* = (sin *θ*)^2^), mean (*μ*), and fixed effects estimates (*e.g., Dr*) the proportional viability of phenotypic marker *Dr^1^* would be:

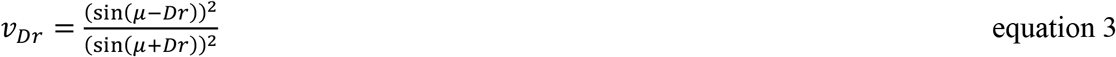

Similarly, the viability-corrected recombination fractions were calculated by estimating the effect of recombinant class from respective ANOVAs, back-transforming, and then pooling across both recombinant classes.

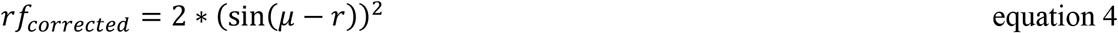

In the results sections both the statistically inferred viability corrected recombination fraction and the raw per bottle recombination fractions (+/− 95% confidence intervals) were shown to illustrate the low repeatability, large uncertainty, and importance of a balanced design that allows statistical correction in recombination experiment.

Finally, one-fourth of X chromosome nondisjunction events in the *F_1_* experimental genotypes are detectable as male white-eyed patroclinous exceptions. The rate of nondisjunction was estimated as four times the number white-eyed males observed per meiosis. Non-disjunction for all arrangements were analyzed jointly after angular transformation with the linear model

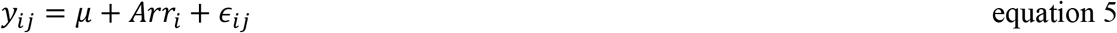

### Testing the Interference Hypothesis

Under the interference hypothesis the recombination intensity will be an increasing function of distance to inversion breakpoint, representing the decay of recombination suppression with increasing genetic distance. This function takes the value zero at the inversion breakpoint indicating fully suppressed recombination, and will increase with a limit of one (wildtype recombination intensity *i.e*. no suppression) in regions most distant from inversion breakpoints. The form of this function and the instantaneous recombination intensity for every intermediate distance is predicted by the coincidence function based on the gamma model of McPeek and Speed (1995) with *D. melanogaster*-specific parameterization (γ = 5, μ = 10). The expected recombination fraction is calculated by integration of this function for the intervals *Gl – Sb*, proximal breakpoint *In(3R)x - Sb*, and distal breakpoint *In(3R)x* – *Dr*. Equations 6A,B give the exact expression for predicted recombination fractions, which are also illustrated in figure 6 and are listed in table 2.

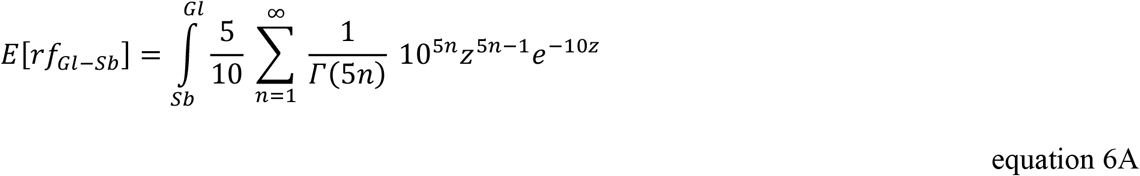

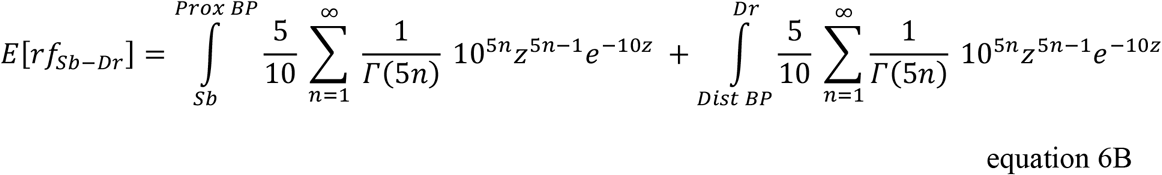

The interference hypothesis was tested with a χ^2^ goodness-of-fit test by comparing the observed count of crossover events corrected for viability effects and the theoretical expectations based on recombination fractions adjusted to sample sizes of the corresponding experiments.

**Figure 5.**
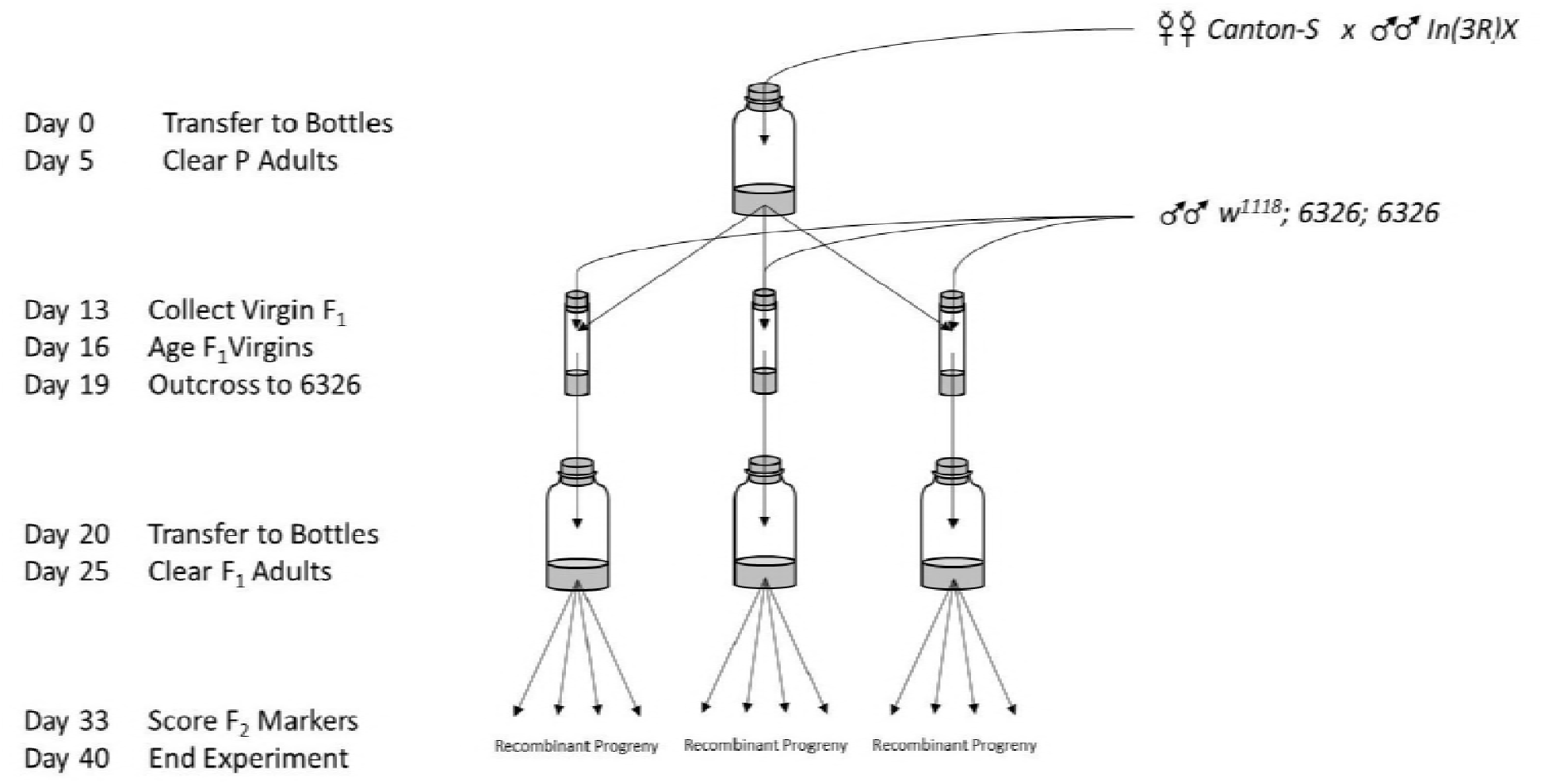
General experimental setup and daily schedule for recombination experiments.

**Figure 6.**
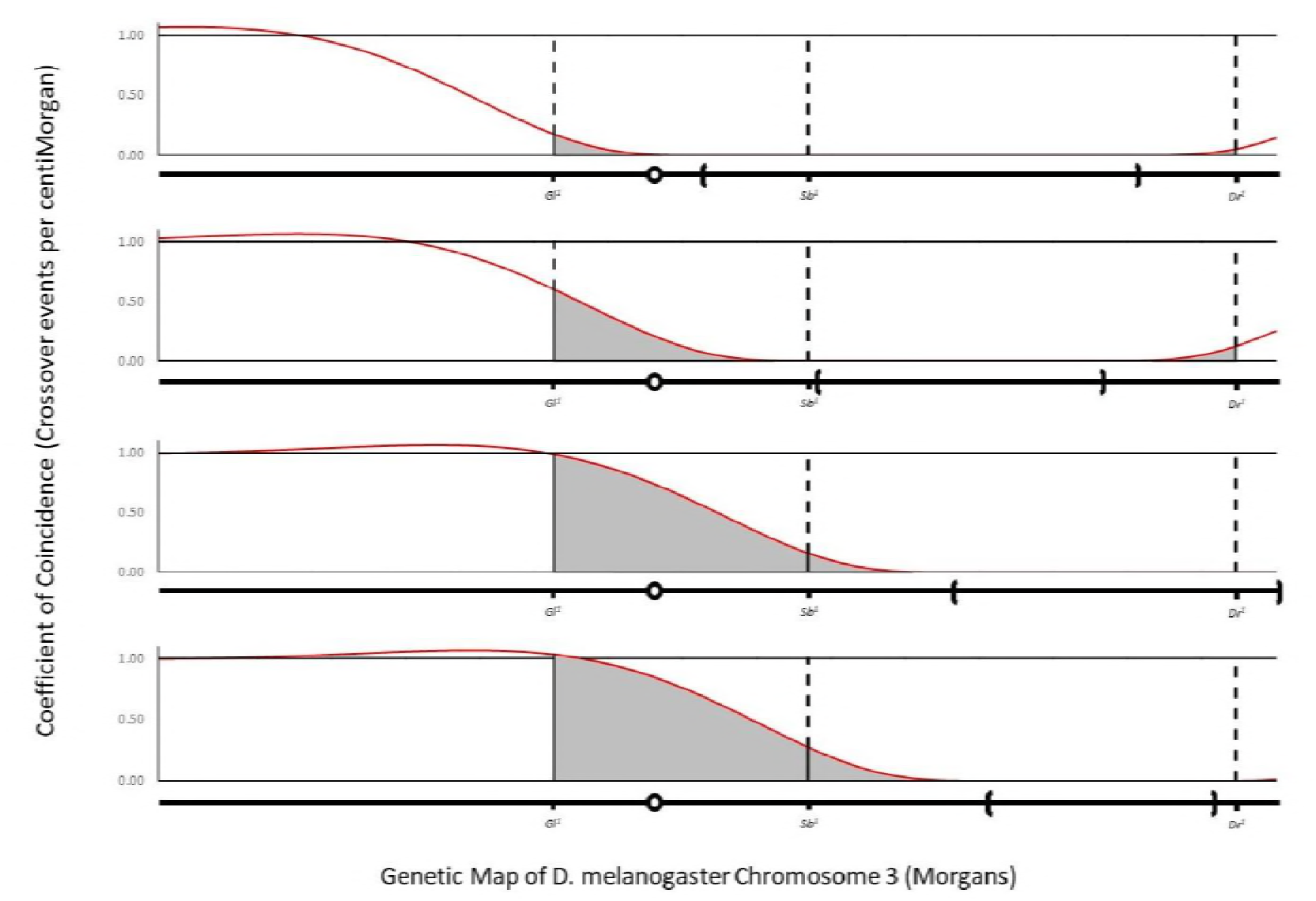
Graphical predictions for the interference hypothesis. Black line is unconditional probability of crossing-over, the red line is the conditional probability for inversion heterozygotes. Expected recombination fraction for a given interval is the gray shaded area under the curve. Position of chromosomal inversions are illustrated below predictions and drawn to scale for the standard genetic map.

**Table 2.**
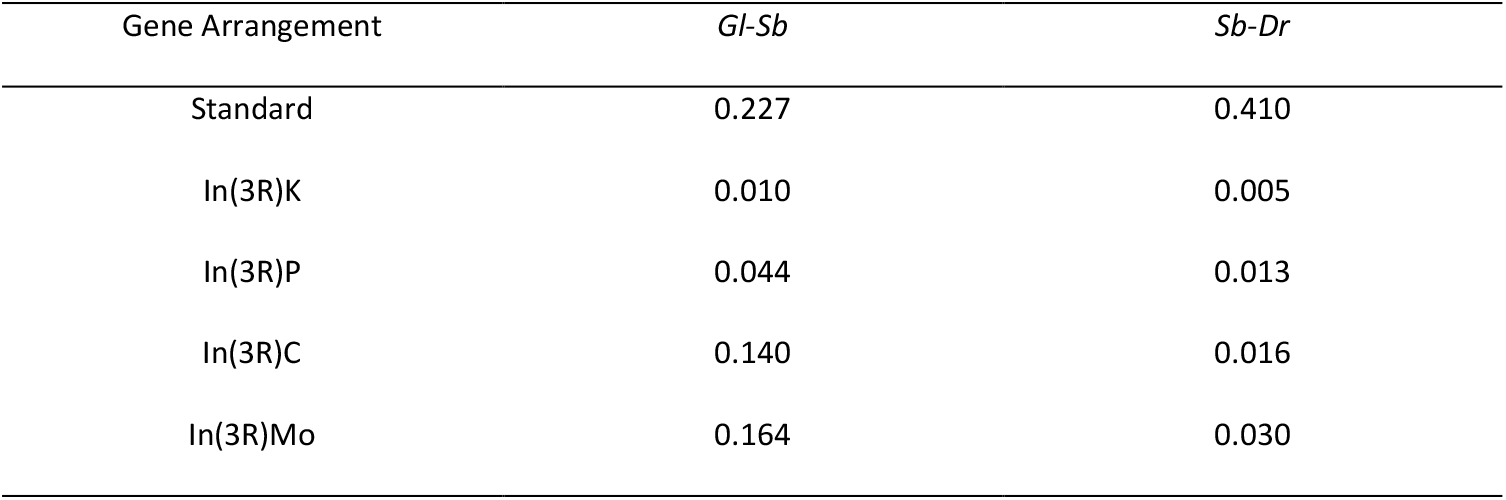
Expected recombination fractions for five alternative third chromosome arrangements based on genetic maps from Comeron *et al*. (2012).

All stocks constructed for these experiments are listed in supplemental table 1 and are available upon request. The author affirms that all data necessary for confirming the conclusions of the article are present within the article, figures, and tables. Full ANOVA tables for every experiment conducted are included as supplemental tables 2–7, and raw count data are available upon request.

## RESULTS

The full experiment scored the product of 44,230 meioses for crossing-over in two intervals for five gene arrangements, consisting of 34 parental crosses and 102 experimental *F_1_* bottles. Each gene arrangement *St, In(3R)C, In(3R)K, In(3R)Mo*, and *In(3R)P* was analyzed separately. Full ANOVA tables for each of the ten experiments (two intervals for five gene arrangements) are included in the supplemental material (supplemental tables 2–6), table 3 in the main text provides a summary of viability main effects, their statistical significance, and the corresponding viability corrected recombination fraction. Under the described experimental conditions the viability-corrected recombination fraction for standard arrangement homokaryotypes interval *Sb* - *Dr* was 0.413 and for the interval *Gl* – *Sb* was 0.186, both of which are consistent with the standard FlyBase genetic map and Comeron *et al*. (2012) estimates of the genetic distances (table 3, supplemental table 2). This result validates the experimental methodology, choice of phenotypic markers, and construction of genetic stocks. Furthermore, it justifies the use of the cytological band-genetic map conversion based on Comeron *et al*. (2012) to generate the expected recombination fraction between inversion breakpoints and phenotypic markers.

**Table 3.** Summary table of the proportional viability of markers or gene arrangement, and the corresponding viability corrected recombination fractions for intervals A) *Sb-Dr* and B) *Gl-Sb*. Asterisks denote statistical significance, confer tables 3-7.

| Arrangement | Stubble | Drop | Inversion | Recombination Fraction |
| --- | --- | --- | --- | --- |
| Standard | 0.935 * | 0.954 | 0.998 | 0.413 |
| In(3R)K | 0.933 * | 1.026 | 0.987 | 1.52E-03 |
| In(3R)P | ----- | 1.017 | 0.996 | 4.49E-03 |
| In(3R)C | 0.911 * | 0.978 | 0.950 | 1.95E-03 |
| In(3R)Mo | 0.943 | ----- | 0.991 | 8.71E-03 |

B
| Arrangement | Glued | Stubble | Inversion | Recombination Fraction |
| --- | --- | --- | --- | --- |
| Standard | 1.013 | 0.812 *** | 0.933 | 0.186 |
| In(3R)K | 0.923 | 0.904 * | 1.178 ** | 0.140 |
| In(3R)P | 0.961 | ----- | 0.932 | 0.113 |
| In(3R)C | 0.888 * | 0.883 * | 0.995 | 0.094 |
| In(3R)Mo | 1.066 * | 0.898 *** | 0.942 | 0.110 |

The rare cosmopolitan inversion *In(3R)K* is the most proximally placed inversion studied and spans 39% of third chromosome standard genetic map. For the interval *Sb* - *Dr*, all 15 observed crossovers are assumed to occur distal to the inversion due to *Sb^1^’s* position internal to inverted region. For the interval *Sb* - *Dr*, accounting for viability effects, back-transforming, and summing over complementary recombinant classes, the viability corrected recombination fraction was estimated as 0.00152 (supplemental table 3A). Between markers *Gl* - *Sb* crossing-over was again assumed to be limited to the proximal region due to *Sb^1^*’s placement inside the inversion. Statistically significant viability effects of *Gl^1^*, the genetic background of *In(3R)K*, and the interaction of these terms were detected (supplemental table 3B). The balanced design allows statistical control of these effects and an estimation of viability-corrected recombination fraction of 0.140.

Common cosmopolitan inversion *In(3R)P* is a medially placed inversion of 26% of the standard genetic map of the third chromosome. The proximal breakpoint of *In(3R)P* is within 4 Mb of *Sb^1^*, disallowing full factorial design (supplemental table 4A). Despite this obstacle, a viability-corrected recombination fraction for interval *Sb* - *Dr* is estimated as 0.00449 by controlling for the effects of *Dr^1^* and the combined effect of *Sb^1^* and the genetic background of *In(3R)P*. A similar procedure based on the partially confounded design for interval *Gl* - *Sb* estimated a viability-corrected recombination fraction of 0.113 (supplemental table 4B).

The rare cosmopolitan inversion *In(3R)C* covers 28% of the standard genetic map and is a terminal inversion (*i.e*., the distal breakpoint is at the telomere). Therefore, marker *Dr^1^* is internal to the inversion and all crossing-over is assumed to occur proximal to the inversion. The viability-corrected recombination fraction of 0.00195 for the interval *Sb* - *Dr* accounts for the statistically significant effects of marker *Sb^1^* (supplemental table 5A). The estimate of viability-corrected recombination fraction for region *Gl* - *Sb* was 0.094 after controlling for statistically significant effects of *Gl^1^, Sb^1^*, and the interaction of *Gl^1^* and genetic background associated with *In(3R)C* (supplemental table 5B).

The smallest and most distally positioned inversion in this study, rare cosmopolitan *In(3R)Mo*, is a subtelomeric rearrangement of 21% of the third chromosome standard genetic map. The distal breakpoint of *In(3R)Mo* is within 5 Mb of marker *Dr^1^*, disallowing a full factorial analysis of interval *Sb* - *Dr* but not affecting the design of experiments for interval *Gl* - *Sb*. The viability-corrected recombination fraction for region between *Sb^1^* and the proximal breakpoint of *In(3R)Mo* is 0.00871 (supplemental table 6A). The viability-corrected fraction for interval *Gl* - *Sb* of 0.110 controls for the statistically significant effects of *Gl^1^, Sb^1^*, and *Gl^1^* * *Sb^1^* (supplemental table 6B)

Analysis of variance for angular transformed rate of non-disjunction revealed no detectable effect of arrangement (*F_4,97_* = 0.537, *p* = 0.709) (supplemental table 7). Non-disjunction rates were on average slightly higher, although without statistical significance, for the standard arrangement homokaryotypes than for chromosomal inversion heterokaryotypes.

A summary of the estimated viability effects and the viability-corrected recombination fractions from all ten experiment are provided in table 3. These are calculated as the fixed effect of marker mutations or inverted gene arrangements extracted from the respective ANOVAs, back-transformed, and then expressed as a proportional reduction in viability assuming wildtype for markers and standard gene arrangement have no viability defects. Inspection of table 3 reveals viability reductions are common (19 of 27), often with statistical significance (10 of 27), but not of uniform magnitude across experiments. Thus, table 3 highlights the importance of correcting for these viability effects in each given experiment estimating recombination rates, the variability due to the linkage array of viability factors for a given cross can be visualize with figure 7C and 7D introduced below. In contrast to the viability effects, the sufficiency of the experimental design and sample size is confirmed by the uniform finding of high statistical significance (*p* << 0.001) for the effect of recombinant versus non-recombinant classes in all experiments (supplemental tables 2–6). To account for viability effects of markers and backgrounds, a corrected recombination rate was estimated by back-transforming the estimated effect of recombinant class from the respective ANOVAs and summing across the two complementary recombinant classes. These corrected recombination rates are reported in the final column of table 3.

**Figure 7.**
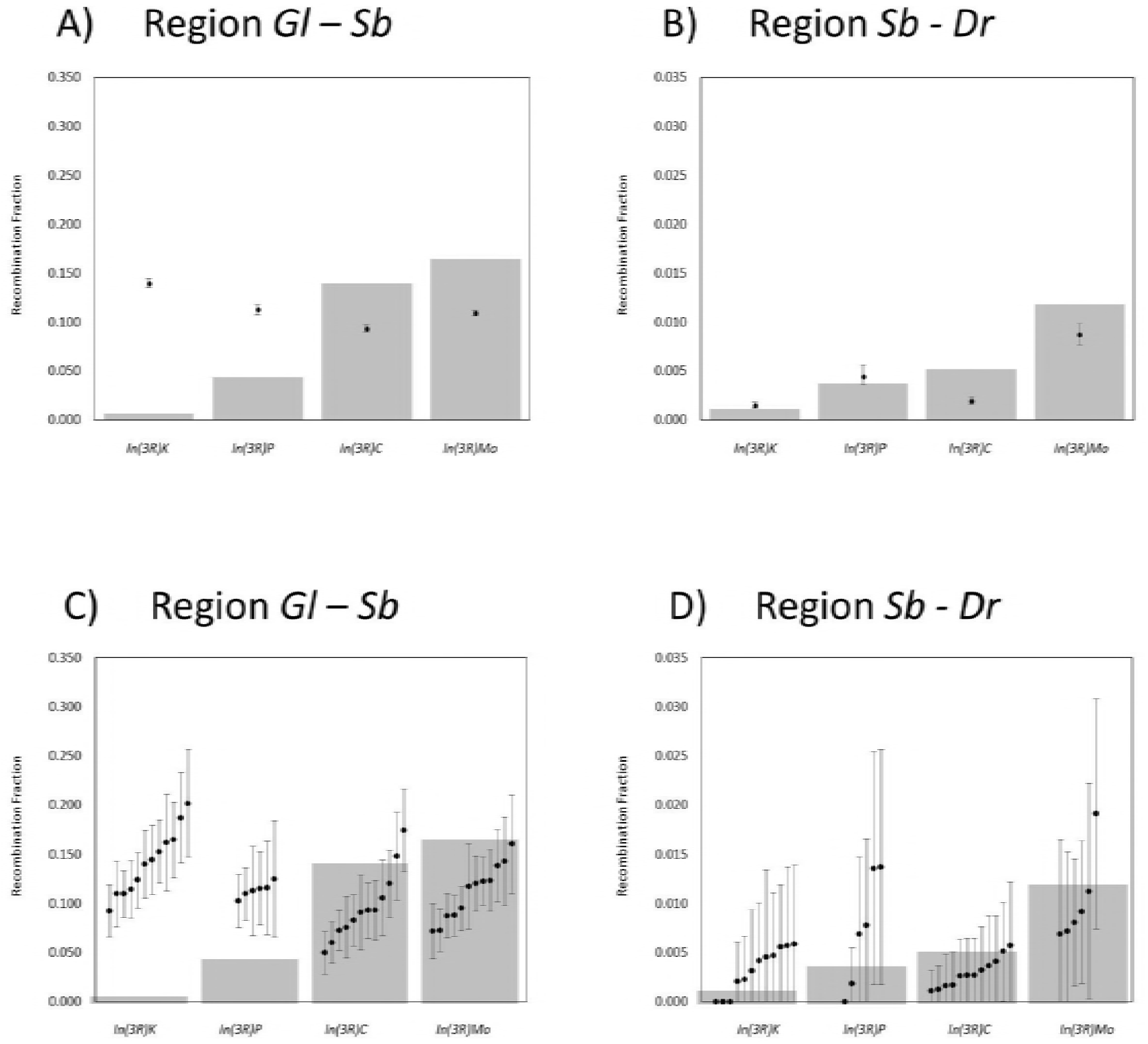
Graphical illustration of the χ^2^ goodness of fit test. Gray bars are expected recombination fractions under the interference hypothesis (equation 6A,B and area under the curve figure 6). Black points are viability corrected recombination fractions +/− one standard error (A and B), and the corresponding raw per bottle recombination fractions with 95% CI (C and D).

Table 4 provides a summary of the goodness-of-fit test including expectations, observations, and viability-corrected estimates of crossover events for experiments on all four cosmopolitan inversions. The predictions of the interference hypothesis were rejected (*χ^2^* = 14621.41, *p* << 0.001). As illustrated in figure 7A and 7B, after viability correction the interference hypothesis under-predicts recombination suppression (*i.e*., observed recombination is less than expected) in both intervals for the two distal inversions *In(3R)C* and *In(3R)Mo*. In contrast, recombination suppression was severely over-predicted (*i.e*., observed recombination is much greater than expected) for the two proximal inversions *In(3R)K* and *In(3R)P*. Approximately 50% of all observed deviation from expectations is attributable to high rates of crossing-over in the centromeric region of the most proximally located inversion *In(3R)K*. The rejection of the interference model due to poor fit in centromere adjacent regions for *In(3R)K* and *In(3R)P* is particularly noticeable upon inspection of the 95% confidence intervals for the recombination fractions calculated on a per experimental bottle basis (figure 7C and 7D).

**Table 4.** Summary table for the *χ*^2^ goodness of fit test of the interference hypothesis including observed, expected, and viability corrected number of crossover events per interval. Asterisks denote statistical significance.

| Arrangement | Interval | Sample Size | Expected | Observed | Viability Corrected | Deviation |
| --- | --- | --- | --- | --- | --- | --- |
| *In(3R)K* | *Sb-Dr* | 4960 | 5.56 | 15 | 7.55 | 0.72 |
| *In(3R)P* | *Sb-Dr* | 2389 | 8.96 | 17 | 10.73 | 0.35 |
| *In(3R)C* | *Sb-Dr* | 8364 | 43.32 | 24 | 16.34 | 16.80 |
| *In(3R)Mo* | *Sb-Dr* | 2985 | 35.40 | 31 | 25.99 | 2.50 |
| *In(3R)K* | *Gl-Sb* | 4774 | 28.73 | 644 | 668.36 | 14242.99 |
| *In(3R)P* | *Gl-Sb* | 1771 | 77.50 | 196 | 200.12 | 194.03 |
| *In(3R)C* | *Gl-Sb* | 4324 | 604.07 | 400 | 404.73 | 65.78 |
| *In(3R)Mo* | *Gl-Sb* | 5463 | 897.93 | 578 | 600.93 | 98.23 |
| | | | | | | $\chi^2_{[7]} = 14621.41$ *** |

## DISCUSSION

Perhaps the least studied effect of chromosomal inversion heterozygosity is the reduction in crossing-over for regions external to inversion breakpoints but still on the same chromosome arm (figure 1A region C). Recombination suppression in this region is enigmatic as it is *not* due to selective elimination of acentric or dicentric chromosomes *nor* is it due to failed pairing, synapsis, or formation of double strand breaks. In the broadest sense, recombination suppression in this region is an interference effect simply because there is an altered probability of crossing-over conditional on structural heterozygosity elsewhere in the genome. Based on cytological analysis, Gong *et al*. (2005) make the more explicit hypothesis that inversion breakpoints possess chiasma-like properties and exert *crossover* interference. Qualitatively, the interference hypothesis is consistent with the general crossover modifying effects observed both internal and external to inverted regions as well as intrachromosomal effects from a wide range of experimental designs and model systems. However, the precise quantitative predictions of this hypothesis systematically fail (*i.e.* consistently under-predicting recombination suppression for distal inversions and consistently over-predicting recombination suppression for proximal inversions) to describe the formal genetic data in the present study.

### Modelling Recombination Suppression

The interference hypothesis is a major development in the study of recombination suppression for inversion heterozygotes because it is the *only* hypothesis to make explicit predictions for the magnitude and extent of recombination suppression external to inverted regions. This is largely based on the pre-existing theoretical treatment of crossover interference, therefore, critical assessment of the current modeling of interference is warranted. There are three possible reasons for the observed deviation from expectations under the interference hypothesis:

1. Incorrect model of interference,
2. Incorrect parameterization of model, or
3. Missing effects in the interference model.

As previously mentioned, recombination suppression external to the inverted region is by definition a form of interference, however, there is no necessary reason the form of this effect must be the same as that for crossover interference. McPeek and Speed (1995) evaluate six alternative models for interference, and although they conclude the gamma model used in the present study is the best mathematical description of crossover interference, it is possible other models may capture the process of recombination suppression more accurately.

Importantly, the form and parameterization of the interference model used in this study was a convergence of several different modeling efforts with different motivations and assumptions (Foss *et al*. 1993; McPeek and Speed 1995; Zhao *et al*. 1995). In addition to *D. melanogaster*, the counting model of Foss *et al*. (1993) has successfully described interference in *Neurosproa crassa, Sacchromyces cerevisiae, Arabidopsis thaliana*, and *Homo sapiens* model systems (Copenhaver *et al*. 2002; Housworth and Stahl 2003; Stahl *et al*. 2004; Berchowitz and Copenhaver 2010, but see Foss and Stahl 1995). The *D. melanogaster* specific parameterization used in this study is derived independently from two gold standard multi-locus recombination datasets (Weinstein 1936; Morgan *et al*. 1938). Notably, the shape parameter inferred for the counting model is in agreement with maximum likelihood estimated from chi-square and gamma models (Foss *et al*. 1993; McPeek and Speed 1995; Zhao *et al*. 1995). Thus, while theoretical developments may improve models and parameterization of crossover interference (*e.g.*, the beam-film model of Kleckner *et al*. 2004; Zhang *et al*. 2014), the explicit predictions of the gamma model of McPeek and Speed (1995) are the appropriate null model for the experimentalist.

Evidence for the third possibility, missing effects in the interference model, can be found in patterns of systematic deviations from expectations. The current interference model overpredicts recombination suppression for *In(3R)K* and *In(3R)P* in both intervals (*i.e*., observed recombination greater than expected), and under-predicts suppression for *In(3R)C* and *In(3R)Mo* (*i.e.*, observed recombination less than expected). There are two major factors unaccounted for in the current model that likely influence interference effects; inversion size and position of inverted regions.

Inversion size would alter conditional probability of crossing-over external to inverted regions only if chiasmata leading to non-crossover gene conversion events in inverted regions exert interference effects, or alternately if there is strong crossover homeostasis. Inversion size does not explain the observed deviations from expectations in these experiments (*r^2^* = 0.39). The relative unimportance of inversion size in the current experiment is consistent with the recent finding that in standard arrangement homokaryotypes non-crossover gene conversion events do not exhibit interference (Miller *et al*. 2016c). Crossover interference and crossover homeostasis have been proposed to be the result of a common patterning process described in the beam-film model (Wang *et al*. 2015). However, crossover homeostasis has not been demonstrated in *D. melanogaster*, and if present is likely to be a weak force in determining the distribution of crossover events (Mehrotra and McKim 2006).

Inversion position along a chromosome would alter conditional probability of crossing-over external to inverted regions if there is a centromere effect on interference. This effect is easily envisioned given the well-known centromere effect on the *unconditional* probability of crossing-over per base pair (Comeron *et al*. 2012; Miller *et al*. 2016c), and the comparative method generalization that distal inversions are stronger suppressors of recombination (Krimbas and Powell 1992). Models of interference, including the gamma model, often assume the effects of interference are uniform with respect to the genetic map (McPeek and Speed 1995). This is the same as assuming either no centromere effect on conditional probabilities scaled to genetic distances, or alternately, centromere effects on conditional and unconditional probabilities are proportional when scaled to physical distance. The observation that recombination suppression is over-predicted for the two proximally placed inversions while under-predicted for the two distally placed inversions, with ~50% of total deviations being attributed to the inversion with breakpoints closest to the centromere, strongly suggests the uniform interference assumption is not valid. Furthermore, under-prediction of recombination suppression by the gamma model for proximal inversions would be expected if conditional probabilities are uniform in physical distance (as in the beam-film model) while unconditional probability of crossing-over per base pair decreases at the centromere. Mathematical evaluation of the hypothesis that crossover interference (conditional probability) scales according to physical distance rather than genetic distance at the centromere is straightforward, but requires higher resolution data than the formal genetic data for two large interval in the present paper.

Confounding the positional effect based on proximity to centromeres is the potential for discontinuities in the decay function introduced by “boundary sites” (Sherizen *et al*. 2005). Formerly thought to be required pairing sites, these chromosomal regions mark a boundary for the recombination suppression effects of translocation heterozygotes used to map their position (Roberts 1970; Roberts 1972; Hawley 1980; Sherizen *et al*. 2005). All four cosmopolitan inversion’s proximal breakpoints lie between the 85A-C and 91A-93D boundary sites and all distal breakpoints lie beyond the last confirmed boundary sites for Muller element E (Sherizen *et al*. 2005). However, the effect of relative proximity of inversion breakpoints to boundary sites within these intervals is unknown. The rough equivalency of observed crossing-over between *Gl* - *Sb* for all four cosmopolitan inversions (figure 7B) could be due to the elimination of interference effects by the 85A-C boundary site centrally placed in this interval. Assuming boundary sites completely inhibit the propagation of interference effects would account 75% of the deviation observed for interval *Gl* - *Sb* of proximally placed inversions *In(3R)K* and *In(3R)P*.

Boundary sites should not be confused with “sensitive sites” (*sensu* Coyne *et al.* 1993) which were not mapped, but chromosomal regions statistically inferred to have the greatest recombination suppression effect based on fertility reduction in pericentric inversion heterozygotes. The proximal breakpoints of *In(3R)C* and *In(3R)Mo* closely coincide with a sensitive site at cytological division 92, which is also the mapped location of a boundary site (*sensu* Sherizen *et al*. 2005). It is, therefore, possible that under-prediction of recombination suppression for the two distal inversions, *In(3R)C* and *In(3R)Mo*, was due to the reduction of the *unconditional* probability of crossing-over for these specific gene arrangements which has not been incorporated into the current predictions. However, under this “sensitive sites” interpretation the over-prediction for distal inversions is unrelated to the under-prediction for the two proximal inversions, which leaves >50% of deviation in these experiments unexplained. Although this discussion highlight a number of paths forward, higher resolution data for a wider range of chromosomal inversions is needed to describe even the most basic patterns of recombination suppression.

### Future Directions

As stated earlier, both crossover interference and recombination suppression by structural heterozygosity are well-known but poorly understood phenomena. The interference hypothesis for recombination suppression in chromosomal inversion heterozygotes and accompanying quantitative predictions establishes the theoretical framework of a null model for testing positional effects associated with boundary sites and centromeres. However, experimental tests manipulating position of inverted regions while rigorously controlling for breakpoint microstructure and genetic background cannot be achieved with X-ray induced rearrangements or inversions drawn from natural populations. Fortunately, development of the *FLP/FRT* recombination system in *D. melanogaster* can be used to synthesize inversions of desired size and position (Golic and Golic 1996). Similarly, advances in whole genome sequencing allows rapid detection of global crossover modifying effects, however, the collection of the all-important rare crossover events in close proximity to inverted regions still presents a logistical problem.

A genetic screen for crossover events adjacent to inversion breakpoints can be conducted by simply following same experimental design described in this study where the *F_1_* virgins with *Sb^1^* and *Dr^1^* in *trans*, are outcrossed to a male double balancer line *w^1118^; 6326; TM3, Sb^1^ e^1^* / *TM6B, Tb^1^ e^1^ Dr^Mio^* (supplemental figure 2). Because the dominant markers used for scoring recombination are lethal recessive and the same as the balancer markers, the result is the selective elimination of 50% of all non-recombinant chromosomes. Viable zygotes with the other 50% of non-recombinant chromosome are easily selected against as they will all be double heterozygotes for *Sb* and *Dr* (either *Dr^1^* / *TM3, Sb^1^ e^1^* or *Sb^1^* / *TM6B, Tb^1^ e^1^ Dr^Mio^*). In addition to elimination of undesirable meiotic products, the viable recombinant chromosome will be marker free (*Sb*^+^ and *Dr*^+^) and automatically preserved in a balanced state in the first generation (heterozygous with either *TM3* or *TM6B* balancer chromosomes).

In the current study, crossovers between dominant phenotypic markers and inversions occurred at an experimentally tractable rate of 10^-2^ to 10^-3^ per meiosis for *Sb^1^* and proximal breakpoints of *In(3R)Mo* or *In(3R)C*. Rates of recombination were similar for *Dr^1^* and the distal breakpoints of *In(3R)P* and *In(3R)K*, but recombination rates below 10^-4^ are indistinguishable from absolute suppression of recombination. In all the preparatory and pilot phases of the experiment, crossing-over was never observed between *Dr^1^* and distal breakpoints of *In(3R)Mo* or *Sb^1^* and the proximal breakpoints of *In(3R)P* out of >10,000 meioses each. Likewise, the double crossover events to introgress *Dr^1^* into *In(3R)C* and *Sb^1^* into *In(3R)K* were only observed once and only after scoring the product of >10,000 meioses combined. Given the logistics of generating these dominant marker-inversion associations *via* a selective screen the proposed genetic screen represents a >75% reduction in effort by eliminating half of non-recombinants and selecting against a single phenotypic class. Employing this method would allow a definitive mapping of the near complete suppression adjacent to inversion breakpoints (figure 1A region A) and provide a means to collect and preserve ample genetic material to fit the true form of the decay with distance function for suppression external to inverted regions (figure 1A region C).

### Conclusion

The discussion highlights a large number of potentially confounding factors, including but not limited to inversion size, inversion position (proximity to boundary sites, sensitive sites, centromeres, and telomeres), breakpoint microstructure, and genetic background. Further complicating matters are the comparative method generalizations based on experiments using heterogeneous live material in terms of genetic backgrounds, inversions, markers, and species. As a result, the magnitude and extent of recombination suppression external to inverted regions is often considered an idiosyncratic behavior of chromosomal inversions, but this lack of understanding is likely due to the absence of mechanistic expectations. The interference hypothesis of recombination suppression for inversions heterozygotes proposed by Gong *et al*. (2005) provides the necessary mechanism to clarify this behavior. Mathematical predictions for this mechanism were generated from a probabilistic model of crossover interference as a stationary renewal point process with gamma distributed interarrival distances. This interference model was tested and rejected based on formal genetic data for crossing-over in two intervals external to inverted regions of four cosmopolitan inversions of *D. melanogaster* Muller element E. Systematic deviation from predictions revealed crossing-over in the centromeric interval of proximally located inversions is much higher than expected. Interestingly, this result suggests the well-known decrease in unconditional probability of crossing-over at centromeres is reversed when considering conditional probabilities. Greater than expected crossing-over in regions near the centromeres when heterozygous for an paracentric inversion can only be detected and evaluated once explicit predictions, such as those derived from the interference hypothesis, are generated for recombination suppression external to inverted regions. Finally, a simple extension of the current experimental system is presented as a genetic screen for the rare crossover events necessary to fit the function describing the decay with distance of recombination suppression external to inverted regions.

## Acknowledgements

This work was designed, conducted, analyzed, and written in partial fulfillment of the author’s Ph.D. dissertation in Ecology and Evolution at Stony Brook University. The work was supported by NIH Project #1R01GM090094-01 awarded to Walt Eanes, the author’s dissertation advisor. The author wishes to thank dissertation committee members Walt Eanes, Josh Rest, and John True of Stony Brook University, as well as David Rand of Brown University, for reading and commenting previous versions of the manuscript. The author also wishes to thank Elise Lauterbur of Stony Brook University for productive discussion on modelling of crossover interference.

---

**Supplemental Table 1.**
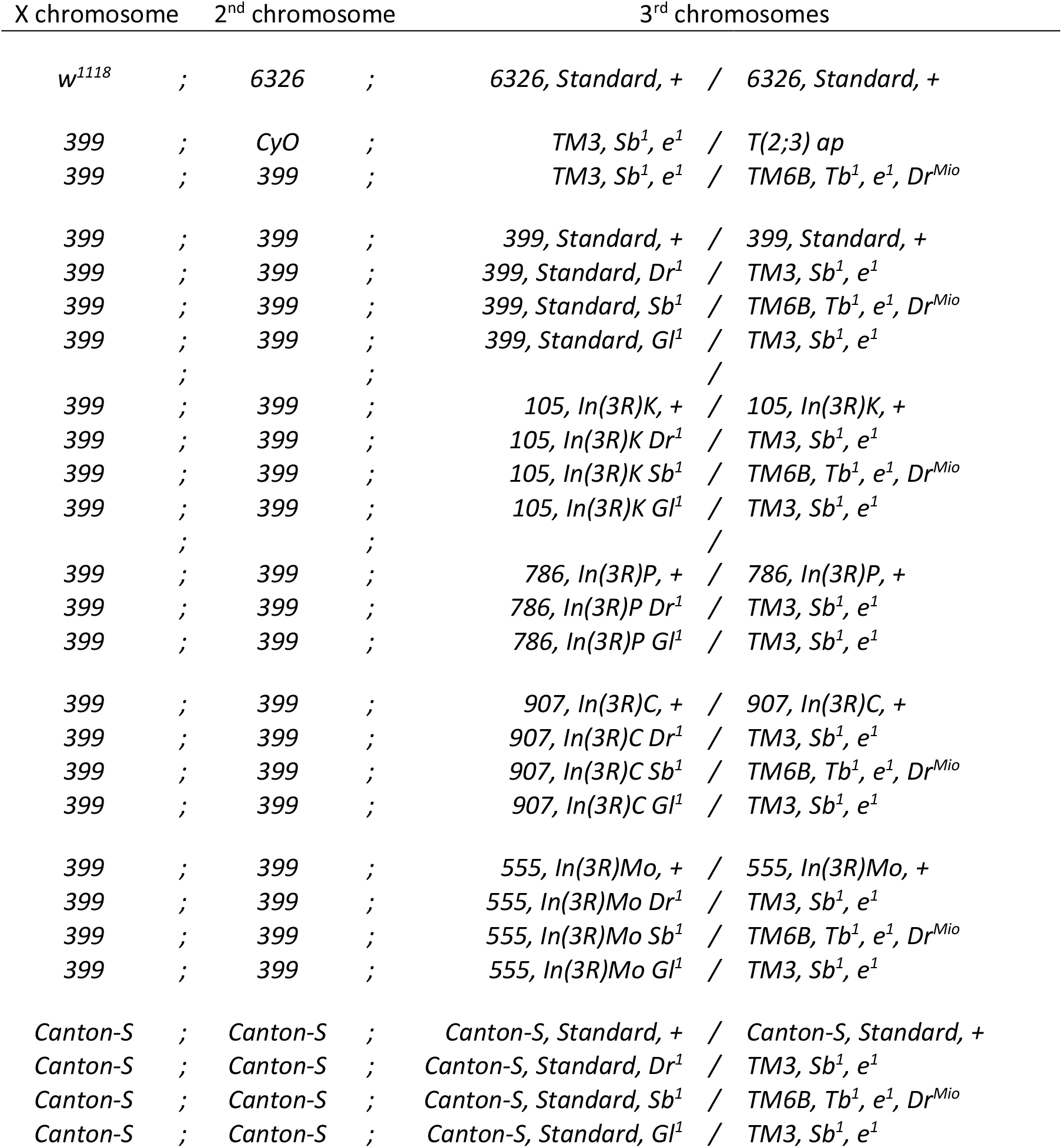
A full list of stocks used in these experiments.

**Supplemental Table 2.** ANOVA table for standard arrangement recombination experiments A) for interval *Sb-Dr* and B) for interval *Gl-Sb*.

| Source of Variation | df | SS | MS | F | p |  |
| --- | --- | --- | --- | --- | --- | --- |
| Recombination | 1 | 397.480 | 397.480 | 197.004 | 0.000 | *** |
| Drop | 1 | 7.307 | 7.307 | 3.622 | 0.064 |  |
| Stubble | 1 | 14.794 | 14.794 | 7.332 | 0.010 | * |
| Arrangement | 1 | 0.011 | 0.011 | 0.005 | 0.942 |  |
| Dr*Sb | 1 | 0.974 | 0.974 | 0.483 | 0.491 |  |
| Dr*Arr | 1 | 0.079 | 0.079 | 0.039 | 0.844 |  |
| Sb*Arr | 1 | 0.006 | 0.006 | 0.003 | 0.957 |  |
| Dr*Sb*Arr | 1 | 1.055 | 1.055 | 0.523 | 0.474 |  |
| Residual Error | 39 | 78.688 | 2.018 |  |  |  |
| Total | 47 | 500.394 |  |  |  |  |

B
| Source of Variation | df | SS | MS | F | p |  |
| --- | --- | --- | --- | --- | --- | --- |
| Recombination | 1 | 5701.128 | 5701.128 | 1244.567 | 0.000 | *** |
| Glued | 1 | 0.466 | 0.466 | 0.102 | 0.751 |  |
| Stubble | 1 | 126.221 | 126.221 | 27.554 | 0.000 | *** |
| Arrangement | 1 | 14.321 | 14.321 | 3.126 | 0.085 |  |
| Gl*Sb | 1 | 4.240 | 4.240 | 0.926 | 0.342 |  |
| Gl*Arr | 1 | 20.506 | 20.506 | 4.477 | 0.041 | * |
| Sb*Arr | 1 | 0.272 | 0.272 | 0.059 | 0.809 |  |
| Gl*Sb*Arr | 1 | 0.097 | 0.097 | 0.021 | 0.885 |  |
| Residual Error | 39 | 178.652 | 4.581 |  |  |  |
| Total | 47 | 6045.903 |  |  |  |  |

**Supplemental Table 3.**
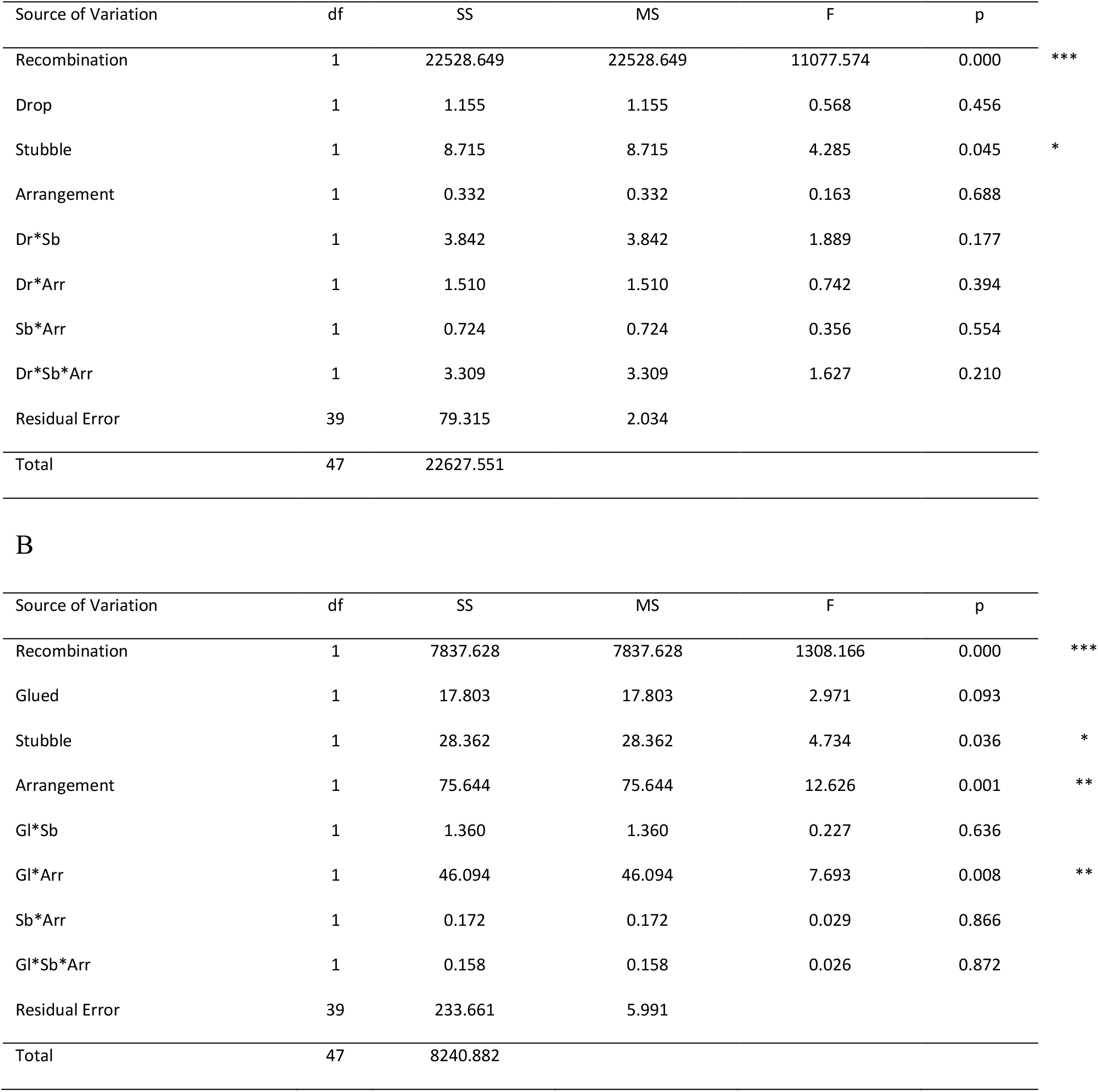
ANOVA table for *In(3R)K* recombination experiments A) *Sb-Dr* and B) *Gl-Sb*.

| Source of Variation | df | SS | MS | F | p |  |
| --- | --- | --- | --- | --- | --- | --- |
| Recombination | 1 | 22528.649 | 22528.649 | 11077.574 | 0.000 | *** |
| Drop | 1 | 1.155 | 1.155 | 0.568 | 0.456 |  |
| Stubble | 1 | 8.715 | 8.715 | 4.285 | 0.045 | * |
| Arrangement | 1 | 0.332 | 0.332 | 0.163 | 0.688 |  |
| Dr*Sb | 1 | 3.842 | 3.842 | 1.889 | 0.177 |  |
| Dr*Arr | 1 | 1.510 | 1.510 | 0.742 | 0.394 |  |
| Sb*Arr | 1 | 0.724 | 0.724 | 0.356 | 0.554 |  |
| Dr*Sb*Arr | 1 | 3.309 | 3.309 | 1.627 | 0.210 |  |
| Residual Error | 39 | 79.315 | 2.034 |  |  |  |
| Total | 47 | 22627.551 |  |  |  |  |

B
| Source of Variation | df | SS | MS | F | p |  |
| --- | --- | --- | --- | --- | --- | --- |
| Recombination | 1 | 7837.628 | 7837.628 | 1308.166 | 0.000 | *** |
| Glued | 1 | 17.803 | 17.803 | 2.971 | 0.093 |  |
| Stubble | 1 | 28.362 | 28.362 | 4.734 | 0.036 | * |
| Arrangement | 1 | 75.644 | 75.644 | 12.626 | 0.001 | ** |
| Gl*Sb | 1 | 1.360 | 1.360 | 0.227 | 0.636 |  |
| Gl*Arr | 1 | 46.094 | 46.094 | 7.693 | 0.008 | ** |
| Sb*Arr | 1 | 0.172 | 0.172 | 0.029 | 0.866 |  |
| Gl*Sb*Arr | 1 | 0.158 | 0.158 | 0.026 | 0.872 |  |
| Residual Error | 39 | 233.661 | 5.991 |  |  |  |
| Total | 47 | 8240.882 |  |  |  |  |

**Supplemental Table 4.** ANOVA table for *In(3R)P* recombination experiments A) *Sb-Dr* and B) *Gl-Sb*.

| Source of Variation | df | SS | MS | F | p |  |
| --- | --- | --- | --- | --- | --- | --- |
| Recombination | 1 | 10620.987 | 10620.987 | 2376.222 | 0.000 | *** |
| Drop | 1 | 0.266 | 0.266 | 0.059 | 0.810 |  |
| Stubble | ---- | ---- | ---- | ---- | ---- |  |
| Arrangement | 1 | 0.013 | 0.013 | 0.003 | 0.958 |  |
| Dr*Sb | ---- | ---- | ---- | ---- | ---- |  |
| Dr*Arr | 1 | 3.752 | 3.752 | 0.839 | 0.371 |  |
| Sb*Arr | ---- | ---- | ---- | ---- | ---- |  |
| Dr*Sb*Arr | ---- | ---- | ---- | ---- | ---- |  |
| Residual Error | 19 | 84.924 | 4.470 |  |  |  |
| Total | 23 | 10709.941 |  |  |  |  |

B
| Source of Variation | df | SS | MS | F | p |  |
| --- | --- | --- | --- | --- | --- | --- |
| Recombination | 1 | 4697.739 | 4697.739 | 979.216 | 0.000 | *** |
| Glued | 1 | 2.169 | 2.169 | 0.452 | 0.509 |  |
| Stubble | ---- | ---- | ---- | ---- | ---- |  |
| Arrangement | 1 | 6.805 | 6.805 | 1.419 | 0.248 |  |
| Gl*Sb | ---- | ---- | ---- | ---- | ---- |  |
| Gl*Arr | 1 | 0.927 | 0.927 | 0.193 | 0.665 |  |
| Sb*Arr | ---- | ---- | ---- | ---- | ---- |  |
| Gl*Sb*Arr | ---- | ---- | ---- | ---- | ---- |  |
| Residual Error | 19 | 91.152 | 4.797 |  |  |  |
| Total | 23 | 4798.791 |  |  |  |  |

**Supplemental Table 5.** ANOVA table for *In(3R)C* recombination experiments A) *Sb-Dr* and B) *Gl-Sb*.

| Source of Variation | df | SS | MS | F | p |  |
| --- | --- | --- | --- | --- | --- | --- |
| Recombination | 1 | 22316.274 | 22316.274 | 7638.438 | 0.000 | *** |
| Drop | 1 | 0.879 | 0.879 | 0.301 | 0.586 |  |
| Stubble | 1 | 15.941 | 15.941 | 5.456 | 0.025 | * |
| Arrangement | 1 | 4.873 | 4.873 | 1.668 | 0.204 |  |
| Dr*Sb | 1 | 0.130 | 0.130 | 0.044 | 0.834 |  |
| Dr*Arr | 1 | 1.874 | 1.874 | 0.641 | 0.428 |  |
| Sb*Arr | 1 | 0.233 | 0.233 | 0.080 | 0.779 |  |
| Dr*Sb*Arr | 1 | 2.704 | 2.704 | 0.926 | 0.342 |  |
| Residual Error | 39 | 113.941 | 2.922 |  |  |  |
| Total | 47 | 22456.849 |  |  |  |  |

B
| Source of Variation | df | SS | MS | F | p |  |
| --- | --- | --- | --- | --- | --- | --- |
| Recombination | 1 | 10584.022 | 10584.022 | 1770.025 | 0.000 | *** |
| Glued | 1 | 37.274 | 37.274 | 6.234 | 0.017 | * |
| Stubble | 1 | 40.744 | 40.744 | 6.814 | 0.013 | * |
| Arrangement | 1 | 0.061 | 0.061 | 0.010 | 0.920 |  |
| Gl*Sb | 1 | 2.778 | 2.778 | 0.465 | 0.500 |  |
| Gl*Arr | 1 | 30.347 | 30.347 | 5.075 | 0.030 | * |
| Sb*Arr | 1 | 7.206 | 7.206 | 1.205 | 0.279 |  |
| Gl*Sb*Arr | 1 | 12.099 | 12.099 | 2.023 | 0.163 |  |
| Residual Error | 39 | 233.204 | 5.980 |  |  |  |
| Total | 47 | 10947.735 |  |  |  |  |

**Supplemental Table 6.** ANOVA table for *In(3R)Mo* recombination experiments A) *Sb-Dr* and B) *Gl-Sb*.

| Source of Variation | df | SS | MS | F | p |  |
| --- | --- | --- | --- | --- | --- | --- |
| Recombination | 1 | 10047.766 | 10047.766 | 3818.301 | 0.000 | *** |
| Drop | ---- | ---- | ---- | ---- | ---- |  |
| Stubble | 1 | 3.475 | 3.475 | 1.321 | 0.265 |  |
| Arrangement | 1 | 0.089 | 0.089 | 0.034 | 0.856 |  |
| Dr*Sb | ---- | ---- | ---- | ---- | ---- |  |
| Dr*Arr | ---- | ---- | ---- | ---- | ---- |  |
| Sb*Arr | 1 | 1.672 | 1.672 | 0.635 | 0.435 |  |
| Dr*Sb*Arr | ---- | ---- | ---- | ---- | ---- |  |
| Residual Error | 19 | 49.998 | 2.631 |  |  |  |
| Total | 23 | 10102.999 |  |  |  |  |

B
| Source of Variation | df | SS | MS | F | p |  |
| --- | --- | --- | --- | --- | --- | --- |
| Recombination | 1 | 9580.159 | 9580.159 | 3989.876 | 0.000 | *** |
| Glued | 1 | 11.003 | 11.003 | 4.582 | 0.039 | * |
| Stubble | 1 | 31.083 | 31.083 | 12.945 | 0.001 | *** |
| Arrangement | 1 | 9.712 | 9.712 | 4.045 | 0.051 |  |
| Gl*Sb | 1 | 21.620 | 21.620 | 9.004 | 0.005 | ** |
| Gl*Arr | 1 | 0.030 | 0.030 | 0.013 | 0.911 |  |
| Sb*Arr | 1 | 2.674 | 2.674 | 1.114 | 0.298 |  |
| Gl*Sb*Arr | 1 | 1.419 | 1.419 | 0.591 | 0.447 |  |
| Residual Error | 39 | 93.644 | 2.401 |  |  |  |
| Total | 47 | 9751.343 |  |  |  |  |

**Supplemental Table 7.**
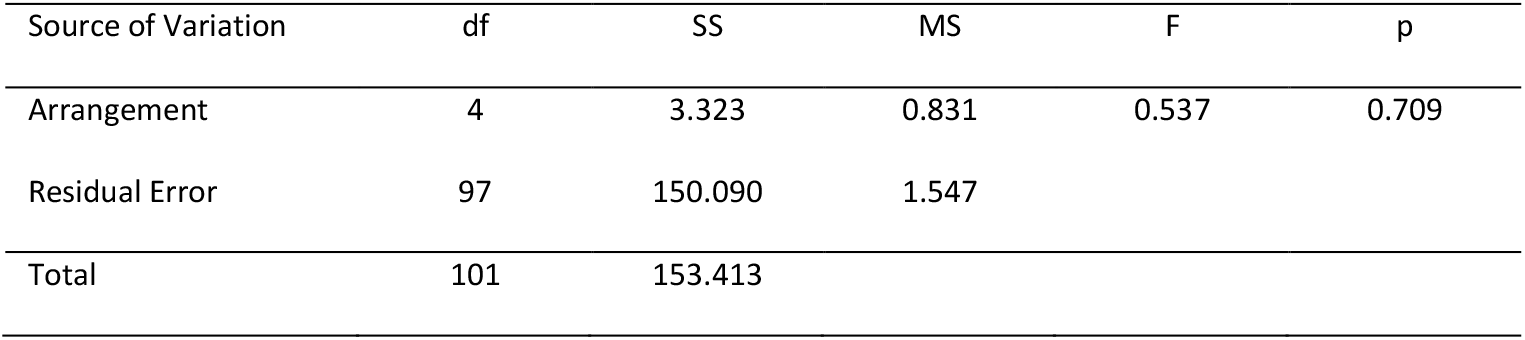
ANOVA table for inferred rates of non-disjunction.

| Source of Variation | df | SS | MS | F | p |
| --- | --- | --- | --- | --- | --- |
| Arrangement | 4 | 3.323 | 0.831 | 0.537 | 0.709 |
| Residual Error | 97 | 150.090 | 1.547 |  |  |
| Total | 101 | 153.413 |  |  |  |

**Supplemental Figure 1.**
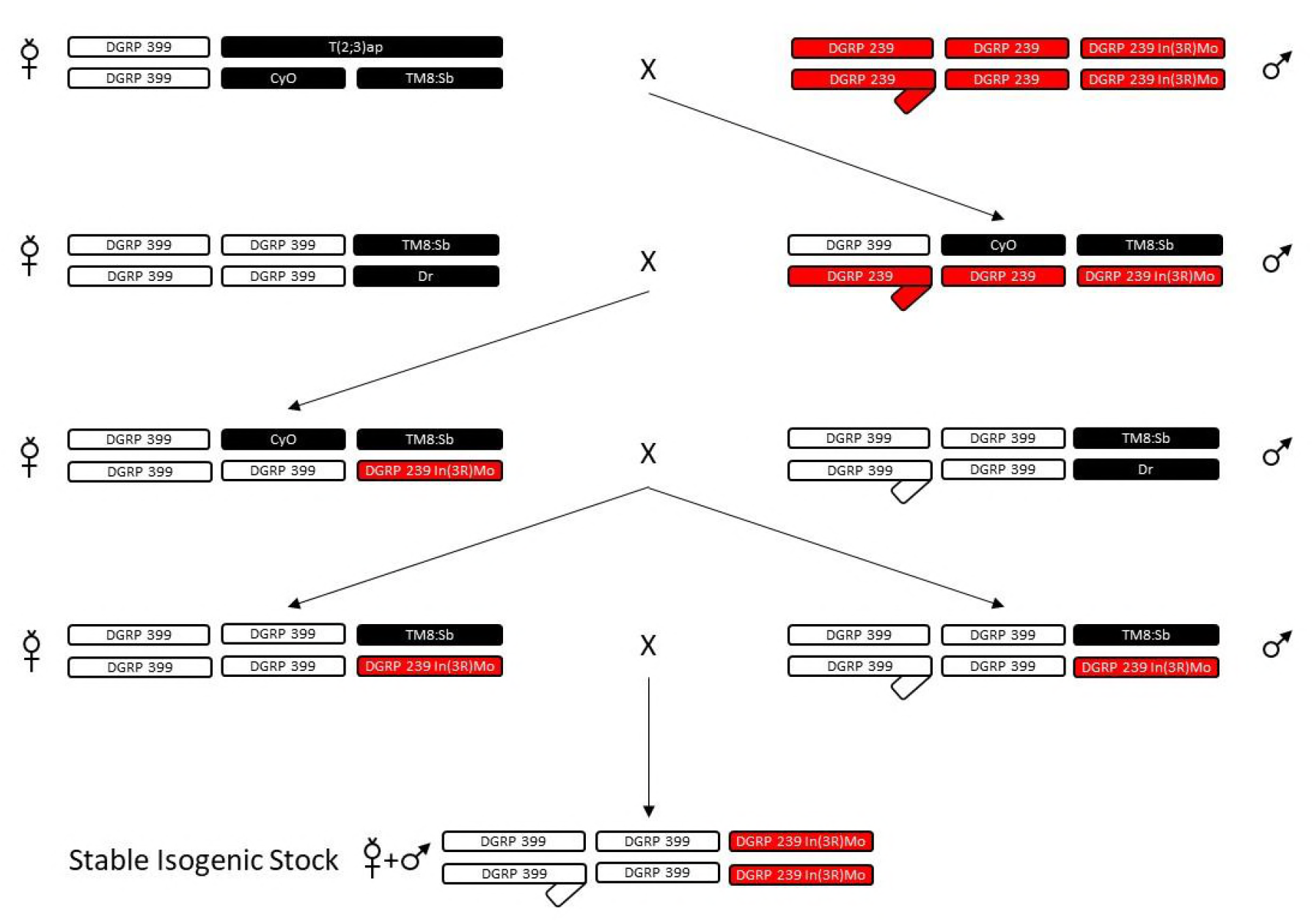
General crossing scheme for background replacement.

**Supplemental Figure 2.**
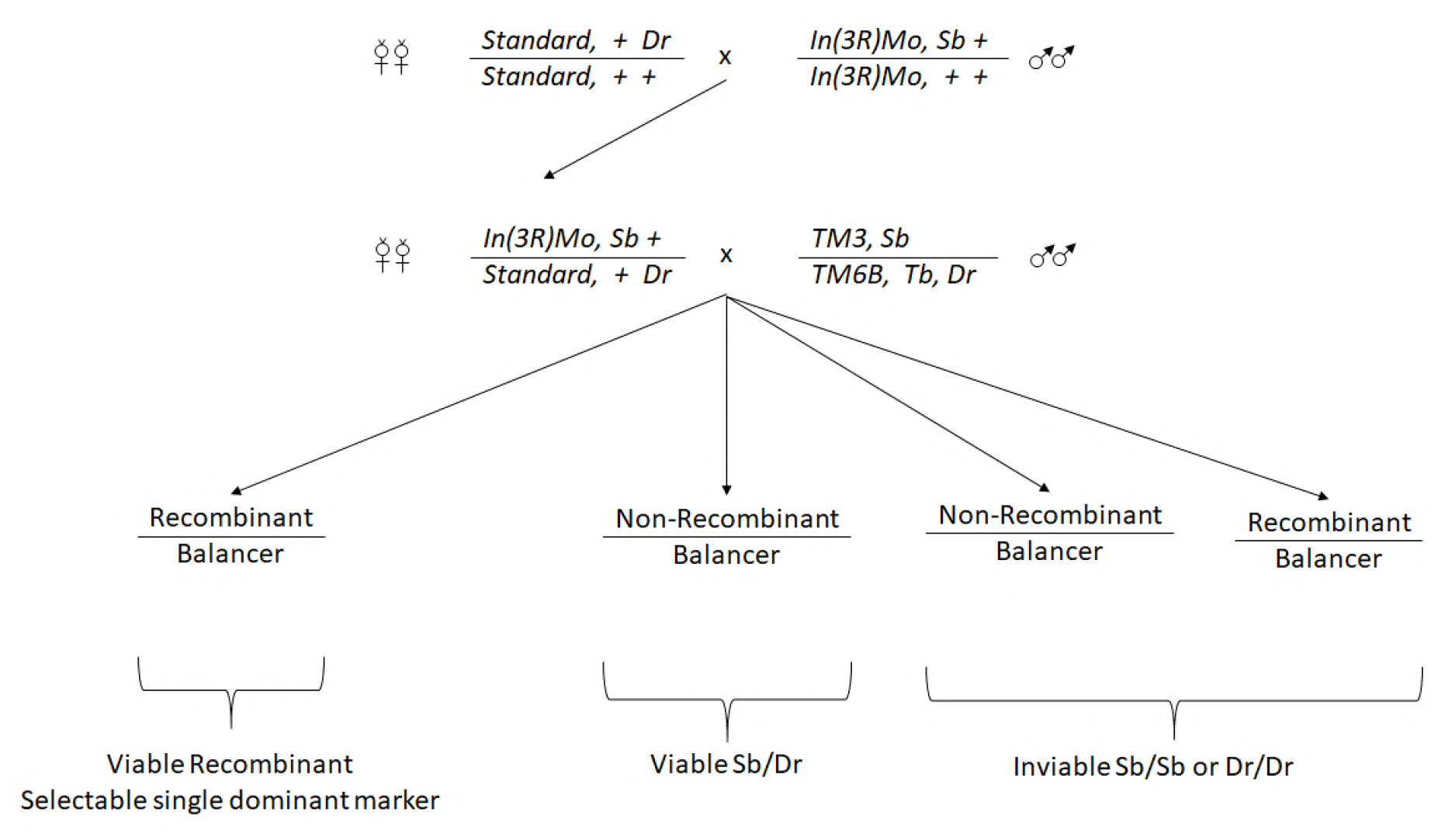
Proposed genetic screen for rare crossovers adjacent to inversions.

## LITERATURE CITED

Bailey, N. T., 1961 Introduction to the mathematical theory of genetic linkage. Claredon Press, Oxford.

Bauer, E., M. Falque, H. Walter, C. Bauland, C. Camisan et al., 2013 Intraspecific variation of recombination rate in maize. Genome Biology 14: R103.

Berchowitz, L. E., and G. P. Copenhaver 2010 Genetic interference: Don’t stand so close to me. Current Genomics 11: 91–102.

Bridges, C. B., and K. S. Brehme, 1944 The mutants of Drosophila melanogaster. Pub. Carnegie Instn. Washington 552: 257.

Carson, H. L., 1946 The selective elimination of inversion dicentric chromatids during meiosis in the eggs of *Sciara impatiens*. Genetics 31: 95–113.

Comeron, J. M., R. Ratnappan and S. Bailin, 2012 The many landscapes of recombination in *Drosophila melanogaster*. PLOS Genetics 8: e1002905.

Copenhaver, G., E. Housworth and F. Stahl, 2002 Crossover interference in *Arabidopsis*. Genetics 160: 1631–1639.

Coyne, J. A., W. Meyers, A. P. Crittenden and P. Sniegowski, 1993 The fertility effects of pericentric inversions in *Drosophila melanogaster*. Genetics 134: 487–496.

Dobzhansky, T., 1931 The decrease of crossing-over observed in translocations, and its probable explanation. The American Naturalist 65: 214–232.

Dobzhansky, T., 1933 Studies on chromosome conjugation. Molecular and General Genetics MGG 64: 269–309.

Dobzhansky, T., and A. Sturtevant, 1931 Translocations between the second and third chromosomes of Drosophila and their bearing on *Oenothera* problems. Pub. Carnegie Instn. Washington 421: 39–59.

Foss, E., R. Lande, F. W. Stahl and C. M. Steinberg, 1993 Chiasma interference as a function of genetic distance. Genetics 133: 681–691.

Foss, E. J., and F. W. Stahl, 1995 A test of a counting model for chiasma interference. Genetics 139: 1201–1209.

Golic, K. G., and M. M. Golic, 1996 Engineering the *Drosophila* genome: chromosome rearrangements by design. Genetics 144: 1693–1711.

Gong, W. J., K. S. McKim and R. S. Hawley, 2005 All paired up with no place to go: pairing, synapsis, and DSB formation in a balancer heterozygote. PLoS Genet 1: e67.

Grell, R. F., 1962a A new model for secondary nondisjunction: the role of distributive pairing. Genetics 47: 1737–1754.

Haldane, J., 1919 The combination of linkage values and the calculation of distances between the loci of linked factors. J Genet 8: 299–309.

Hawley, R. S., 1980 Chromosomal sites necessary for normal levels of meiotic recombination in *Drosophila melanogaster*. I. Evidence for and mapping of the sites. Genetics 94: 625–646.

Hoffmann, A. A., and L. H. Rieseberg, 2008 Revisiting the impact of inversions in evolution: from population genetic markers to drivers of adaptive shifts and speciation? Annual Review of Ecology, Evolution, and Systematics 39: 21–42.

Housworth, E., and F. Stahl, 2003 Crossover interference in humans. The American Journal of Human Genetics 73: 188–197.

Hunter, C. M., W. Huang, T. F. Mackay and N. D. Singh, 2016 The genetic architecture of natural variation in recombination rate in *Drosophila melanogaster*. PLoS Genet 12: e1005951.

Johnston, S. E., C. Bérénos, J. Slate and J. M. Pemberton, 2016 Conserved genetic architecture underlying individual recombination rate variation in a wild population of Soay sheep (*Ovis aries*). Genetics 203: 583–598.

Kirkpatrick, M., and N. Barton, 2006 Chromosome inversions, local adaptation and speciation. Genetics 173: 419–434.

Kleckner, N., D. Zickler, G. H. Jones, J. Dekker, R. Padmore et al., 2004 A mechanical basis for chromosome function. Proc. Nat. Acad. Sci. 101: 12592–12597.

Kong, A., G. Thorleifsson, M. L. Frigge, G. Masson, D. F. Gudbjartsson et al., 2014 Common and low-frequency variants associated with genome-wide recombination rate. Nat Genet 46: 11–16.

Kosambi, D. D., 1943 The estimation of map distances from recombination values. Annals of Eugenics 12: 172–175.

Krimbas, C. B., and J. R. Powell, 1992 Introduction, pp. 1–52 in Drosophila inversion polymorphism, edited by C. B. Krimbas and J. R. Powell. CRC Press, Boca Raton, FL.

Langley, C. H., K. Stevens, C. Cardeno, Y. C. G. Lee, D. R. Schrider et al., 2012 Genomic variation in natural populations of *Drosophila melanogaster*. Genetics: 192: 533–598.

Lucchesi, J. C., and D. T. Suzuki, 1968 The interchromosomal control of recombination. Annual Review of Genetics 2: 53–86.

Mackay, T. F. C., S. Richards, E. A. Stone, A. Barbadilla, J. F. Ayroles et al., 2012 The *Drosophila melanogaster* genetic reference panel. Nature 482: 173–178.

Madan, K., M. Seabright, R. Lindenbaum and M. Bobrow, 1984 Paracentric inversions in man. Journal of Medical Genetics 21: 407–412.

McClintock, B., 1941 The stability of broken ends of chromosomes in *Zea mays*. Genetics 26: 234–282.

McPeek, M. S., and T. P. Speed, 1995 Modeling interference in genetic recombination. Genetics 139: 1031–1044.

Mehrotra, S., and K. McKim, 2006 Temporal analysis of meiotic DNA double-strand break formation and repair in *Drosophila* females. PLoS Genet 2: e200.

Miller, D. E., K. R. Cook, A. V. Arvanitakis and R. S. Hawley, 2016a Third chromosome balancer inversions disrupt protein-coding genes and influence distal recombination events in *Drosophila melanogaster*. G3: Genes| Genomes| Genetics 6: 1959–1967.

Miller, D. E., K. R. Cook, N. Y. Kazemi, C. B. Smith, A. J. Cockrell et al., 2016b Rare recombination events generate sequence diversity among balancer chromosomes in *Drosophila melanogaster*. Proc. Nat. Acad. Sci. 113: E1352–E1361.

Miller, D. E., C. B. Smith, N. Y. Kazemi, A. J. Cockrell, A. V. Arvanitakis et al., 2016c Whole-genome analysis of individual meiotic events in *Drosophila melanogaster* reveals that noncrossover gene conversions are insensitive to interference and the centromere effect. Genetics 203: 159–171.

Morgan, T. H., C. Bridges and J. Schultz, 1938 Constitution of the germinal material in relation to heredity. Pub. Carnegie Instn. Washington 37: 304–309.

Muller, H. J., 1916 The mechanism of crossing-over. The American Naturalist 50: 193–221.

Navarro, A., and A. Ruiz, 1997 On the fertility effects of pericentric inversions. Genetics 147: 931–933.

Novitski, E., and G. Braver, 1954 An analysis of crossing over within a heterozygous inversion in *Drosophila melanogaster*. Genetics 39: 197–209.

Painter, T. S., 1933 A new method for the study of chromosome rearrangements and the plotting of chromosome maps. Science 78: 585–586.

Roberts, P., 1972 Differences in synaptic affinity of chromosome arms of *Drosophila melanogaster* revealed by differential sensitivity to translocation heterozygosity. Genetics 71: 401–415.

Roberts, P. A., 1970 Screening for x-ray-induced crossover suppressors in *Drosophila melanogaster*: prevalence and effectiveness of translocations. Genetics 65: 429–448.

Schultz, J., and H. Redfield, 1951 Interchromosomal effects on crossing over in *Drosophila*, pp. 175–197 in Cold Spring Harbor symposia on quantitative biology. Cold Spring Harbor Laboratory Press.

Sherizen, D., J. K. Jang, R. Bhagat, N. Kato and K. S. McKim, 2005 Meiotic recombination in *Drosophila* females depends on chromosome continuity between genetically defined boundaries. Genetics 169: 767–787.

Sokal, R. R., and F. J. Rohlf, 1995 Biometry: the principles and practice of statistics in biological research. W. H. Freeman and Company, New York.

Speed, T. P., 1996 What is a genetic map function?, pp. 65–88 in Genetic Mapping and DNA sequencing edited by T. Speed and M. Waterman. Springer, New York.

Stahl, F. W., H. M. Foss, L. S. Young, R. H. Borts, M. F. Abdullah et al., 2004 Does crossover interference count in *Saccharomyces cerevisiae*? Genetics 168: 35–48.

Stevison, L. S., K. B. Hoehn and M. A. F. Noor, 2011 Effects of inversions on within- and between-species recombination and divergence. Genome Biology and Evolution 3: 830–841.

Sturtevant, A., 1920 Genetic studies on *Drosophila simulans*. I. Introduction. Hybrids with *Drosophila melanogaster*. Genetics 5: 488–500.

Sturtevant, A., 1921 A case of rearrangement of genes in *Drosophila*. Proc. Nat. Acad. Sci. 7: 235–237.

Sturtevant, A., 1931 Known and probable inverted sections of the autosomes of *Drosophila melanogaster*. Pub. Carnegie Instn 421: 1–27.

Sturtevant, A., and E. Novitski, 1941 The homologies of the chromosome elements in the genus *Drosophila*. Genetics 26: 517–541.

Sturtevant, A. H., 1913 The linear arrangement of six sex-linked factors in *Drosophila*, as shown by their mode of association. Journal of Exper. Zoo. A: Ecol. Genet. and Phys. 14: 43–59.

Sturtevant, A. H., 1915 The behavior of the chromosomes as studied through linkage. Molecular and General Genetics MGG 13: 234–287.

Sturtevant, A. H., 1917 Genetic factors affecting the strength of linkage in *Drosophila*. Proc. Natl. Acad. Sci. 3: 555–558.

Sturtevant, A. H., and G. W. Beadle, 1936 The relations of inversions in the X chromosome of *Drosophila melanogaster* to crossing over and disjunction. Genetics 21: 554–604.

Sturtevant, A. H., and T. Dobzhansky, 1936 Inversions in the third chromosome of wild races of *Drosophila pseudoobscura*, and their use in the study of the history of the species. Proc. Natl. Acad. Sci. 22: 448–450.

Sturtevant, A. H., and K. Mather, 1938 The interrelations of inversions, heterosis and recombination. The American Naturalist 72: 447–452.

Wang, S., D. Zickler, N. Kleckner and L. Zhang, 2015 Meiotic crossover patterns: obligatory crossover, interference and homeostasis in a single process. Cell Cycle 14: 305–314.

Weinstein, A., 1936 The theory of multiple-strand crossing over. Genetics 21: 155–199.

Zhang, L., Z. Liang, J. Hutchinson and N. Kleckner, 2014 Crossover patterning by the beam-film model: analysis and implications. PLoS Genet 10: e1004042.

Zhao, H., T. P. Speed and M. S. McPeek, 1995 Statistical analysis of crossover interference using the chi-square model. Genetics 139: 1045–1056.

